# Peptide inhibitors recognize prefusion viral fusion proteins with heterogeneous stoichiometry and rapid kinetics

**DOI:** 10.64898/2026.09.14.750718

**Authors:** Revansiddha H. Katte, Junyu Liu, Yufan He, Bhishem Thakur, Xiao Huang, Jared J. Lindenberger, Wang Xu, Narendra Kumar Gonepudi, Yang Han, Katarzyna Janowska, Robert J. Edwards, Priyamvada Acharya, Maolin Lu

## Abstract

Peptide fusion inhibitors, an important class of antivirals, block viral entry by targeting fusion proteins required for membrane fusion. However, their interactions with intact trimeric fusion proteins remain elusive; direct observation of binding on virions or native-like trimers has been lacking. Here, we developed a single-molecule imaging platform to visualize peptide binding in real time. LP-98 bound HIV-1 Envelope (Env) trimers on virions and, unexpectedly, prefusion-stabilized soluble Env trimers, with higher affinity for virion-associated Env and, among soluble trimers, a mutant Env. RSV fusion-inhibiting T-118 and 4ca similarly engaged prefusion-stabilized fusion (F) trimers with rapid kinetics and high-nanomolar affinities. Binding to prefusion Env or F demonstrates that peptide inhibitors can act earlier than the canonical prehairpin-intermediate model suggests. Individual binding events revealed heterogeneous peptide-to-trimer stoichiometries, with single-peptide occupancy predominating, while stepwise and simultaneous events revealed multiple routes to higher occupancy. These findings expand the canonical model of peptide fusion inhibition and provide previously inaccessible mechanistic insights into how antiviral peptide fusion inhibitors act.

## Introduction

Enveloped viruses rely on fusion proteins to mediate membrane fusion for cellular entry. Class I fusion proteins, employed by viruses such as human immunodeficiency virus type 1 (HIV-1), influenza virus, severe acute respiratory syndrome coronavirus 2 (SARS-CoV-2), and respiratory syncytial virus (RSV), are metastable^1^. They undergo large conformational rearrangements from pre- to post-fusion states and represent important targets for antiviral intervention^1–7^. These viral fusion proteins contain two complementary heptad-repeat (HR) helical regions, termed HR1 and HR2 in HIV-1 Envelope (Env) and HRA and HRB in RSV fusion (F) protein^8–11^. During membrane fusion, these regions associate to form the six-helix coiled-coil bundle (6HB) that drives membrane apposition^8–11^. This conserved mechanism underpins heptad-repeat-mimicking peptide fusion inhibitors, an important class of antiviral therapeutics^12,13^. They mimic the C-terminal HR2 of HIV-1 Env, the HRB region of RSV F, or equivalent region of other class I fusion proteins and bind the complementary HR1/HRA region exposed during fusion protein refolding^14–20^, thus preventing formation of the 6HB required for membrane fusion^1,2,21,22^.

Given the high conservation of these complementary target regions, these inhibitors can retain broad activity across diverse viral strains and subtypes. Mechanistically, according to the prevailing model, their inhibitory interaction with fusion proteins occurs after activation of the prefusion trimer, when the N-terminal heptad-repeat region becomes exposed in the transient putative prehairpin intermediate^2,10,11,21–23^. However, much of the structural and biochemical evidence supporting this model comes from studies using fusion-protein-derived peptide fragments^11,20,24–26^, leaving open the question of whether they can engage native or native-like trimers before formation of this intermediate. Thus, despite extensive characterization of their antiviral potency, breadth, and proposed mechanisms of action, how peptide fusion inhibitors directly interact with trimeric or native membrane-associated fusion proteins remains poorly defined, particularly with respect to fusion stage, binding kinetics, stoichiometry, and molecular heterogeneity.

Here, we developed a real-time single-molecule imaging approach that resolves binding kinetics, stoichiometry, sequential versus simultaneous binding modes, and event-to-event heterogeneity. Using this approach, we characterized binding of LP-98^16^, a recently developed HIV-1 HR2-derived peptide fusion inhibitor, to native Env on intact virions and in native-like soluble trimeric Env ectodomains. We also characterized the binding of RSV HRB-derived peptide inhibitors T-118^18^ and 4ca^19^ to the fusion (F) protein trimers. Across both viral systems, peptide inhibitors engaged prefusion-stabilized Env and F trimers, demonstrating that their direct interaction is not restricted to the long-proposed putative prehairpin intermediate. Single-molecule analysis further revealed association and dissociation kinetics, heterogeneous binding stoichiometries, and simultaneous and sequential binding events. These findings expand the mechanistic view of how peptide fusion inhibitors engage dynamic viral fusion proteins.

## Results

### Establishing a single-molecule ligand-binding imaging strategy

We first established a single-molecule imaging platform based on a customized prism-based total internal reflection (TIRF) fluorescence microscope (**Fig. 1A**), capable of imaging intact virions (**Fig.1B**) and recombinant proteins and of resolving binding kinetics and stoichiometry (**Fig.1**). In this strategy, the ligand and target molecule are labeled with spectrally distinct fluorophores, enabling their interactions to be tracked in real time (**Fig. 1C**). We leveraged validated labeling sites (**Figs. S1-S2**), labeling approaches, and imaging strategies from our previous single-molecule FRET studies^27–30^ and applied them here. For Env on intact HIV-1 virions, we site-specifically labeled Env with a Cy5-derived fluorophore using previously established protocols involving genetic code expansion and click chemistry^29,31,32^. For soluble Env or F trimers, we expressed and purified proteins carrying a short peptide tag (**Figs. S1** and **S3**) for site-specific enzymatic labeling with a Cy5-derived fluorophore (see **Methods**), as previously described^27^.

**Figure 1.**
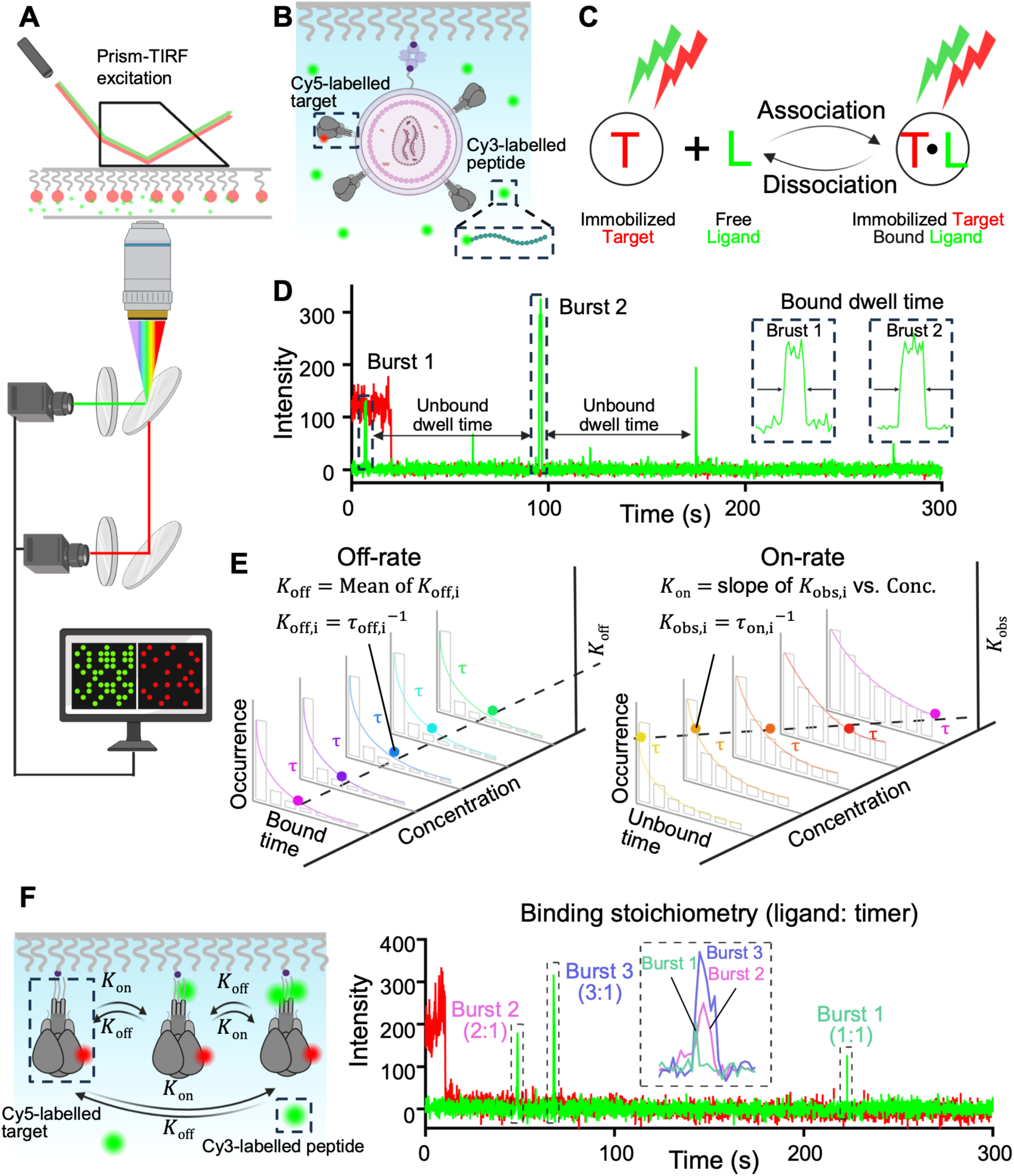
Real-time single-molecule imaging platform for quantifying ligand–target binding kinetics on virions and with recombinant targets. **(A)** Single-molecule TIRF microscopy setup for monitoring real-time target-ligand binding. **(B)** Schematic of binding partners: an immobilized intact virion carrying a Cy5-labeled target (e.g., HIV-1 Env) and freely diffusing Cy3-labeled ligands (e.g., peptides). **(C)** Binding events are identified by the simultaneous detection of colocalized Cy5 and Cy3 fluorescence signals, indicating formation of a target-ligand complex. **(D)** Representative fluorescence trajectory showing association and dissociation of Cy3-labeled peptides with Cy5-labeled HIV-1 Env. Green fluorescence bursts indicate individual binding events. Bound dwell time is defined as the duration of each burst, whereas unbound dwell time is the interval between consecutive binding events. **(E)** Estimation of kinetic parameters from single-molecule trajectories. Left: The dissociation rate constant (off-rate) is determined from the distribution of bound dwell times. Right: The association rate constant (on-rate) is obtained from the concentration-dependent slope of rates derived from unbound dwell time distributions. **(F)** Direct quantification of binding stoichiometry from individual trajectories. A trimeric recombinant target protein (e.g., HIV-1 Env or RSV F) can bind up to three peptide ligands. The number of bound ligands (1:1, 2:1, or 3:1) is determined from the fluorescence intensity associated with each binding event (left, schematic; right, representative experimental trace).

To demonstrate this approach (**Figs.1B-E**), labeled HIV-1 virions were immobilized on a quartz surface in a stop-flow reaction chamber and incubated with a defined concentration of Cy3-labeled Env HR2-derived peptide inhibitor LP-98^16^. Both fluorophores were simultaneously excited, and their fluorescence signals were split by a beamsplitter and recorded synchronously with two separate cameras (see **Methods**). Fluorescence intensity trajectories were extracted from the Cy3 and Cy5 channels, from which individual binding events were derived (**Fig. 1D**). Binding events were identified as bursts of Cy3 fluorescence that colocalized with Cy5-labeled virions (**Fig. 1D**). With Cy5 fluorescence marking virion positions, binding events were analyzed from the Cy3 trajectories, in which fluorescence bursts and baseline periods represented the bound and unbound states, respectively. We defined the bound dwell time as the full width at half-maximum of individual fluorescence bursts, and the unbound dwell time as the time interval between two consecutive bursts (**Fig.1D**).

To quantify binding kinetics, we titrated fusion protein with ligand over a range of concentrations and compiled the resulting bound and unbound dwell-time histograms at each concentration (*i*) (**Fig. 1E**). The distributions were fitted with single-exponential decay functions to obtain the corresponding time constants (*τ*_on,i_ and *τ*_off,i_) (**Fig.1E**). *τ*_off,i_ derived from bound dwell times were used to calculate the dissociation rate constant (*K*_off,i_), whereas *τ*_on,i_ derived from unbound dwell times yielded the observed association rate constants ( *K*_obs,i_). Because *K*_off_ is independent of ligand concentration, values obtained across all concentrations were averaged to determine the final *K*_off_. In contrast, *K*_obs,i_ increased as a function of ligand concentration. Plotting *K*_obs,i_ as a function of ligand concentration enabled determination of the association rate constant (*K*_on_), from which the equilibrium dissociation constant (*K*_d_) was calculated (**Fig. 1E**). Detailed equations and fitting procedures are described in the Methods section (**Eqs. 1 – 8**)^33,34^.

The same imaging and analysis strategy was applied to soluble proteins, as demonstrated with Cy5-labeled soluble Env trimers and Cy3-labeled LP-98 (**Fig. 1F**). Beyond binding kinetics, individual fluorescence trajectories enabled peptide-to-trimer binding stoichiometry to be determined from the fold increase in Cy3 fluorescence intensity relative to the mean intensity of a single Cy3-labeled peptide.

### LP-98 binding kinetics to native Env trimers on intact HIV-1 virions

We next applied our established single-molecule approach to directly quantify LP-98 binding to Env on intact HIV-1 virions (**Figs. 2, S4, and S5**). We examined two primary HIV-1 Env isolates: BG505 (full-length HIV-1_BG505_ virions, **Figs. 2 and S4**) and JR-FL (HIV-1 particles pseudotyped with JR-FL Env - HIV-1_JR-FL_, **Fig.S5**). LP-98^16^ is an HIV-1 peptide fusion inhibitor derived from the gp41 HR2 region of Env (the high-resolution structure is simulated^35^) (**Fig. 2A**). During HIV-1 entry, Env undergoes conformational rearrangements that culminate in the formation of the gp41 6HB, in which three HR2 helices pack against a trimeric HR1 core^36^ (**Fig. 2B**). By mimicking HR2, LP-98 is thought to inhibit fusion by competing with HR2 for binding with HR1 during 6-HB formation^26^. Despite its exceptional antiviral potency (**Fig. 2C**, sub-picomolar IC50 tested in our hands) and promising preclinical performance^16,37^, the binding kinetics of LP-98 to Env have not been directly characterized.

**Figure 2.**
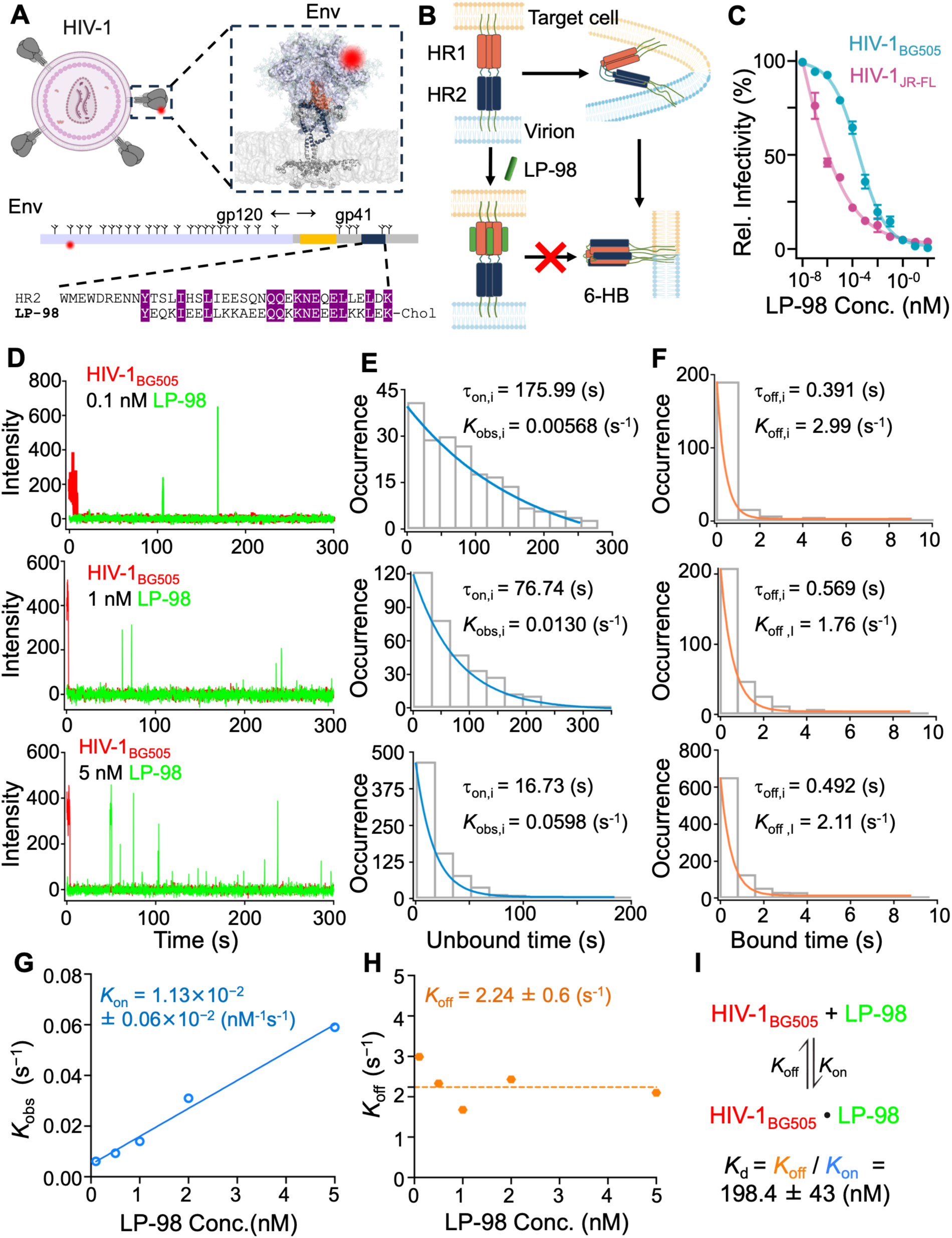
Quantification of LP-98 binding kinetics to native Env_BG505_ trimers on intact virions, HIV-1_BG505_, from individual events. **(A)** Schematic of virion-associated HIV-1 Env trimer labeled by a Cy5-derived fluorophore (red). Env is composed of gp120 and gp41 subunits. The gp41 heptad-repeat regions HR1 (yellow) and HR2 (navy blue) are critical for membrane fusion. LP-98 is an HR2-derived fusion-inhibitory peptide, with conserved residues highlighted. **(B)** The prevailing model of LP-98-mediated fusion inhibition. LP-98 binds to HR1, preventing formation of the postfusion 6HB and thereby blocking membrane fusion. **(C)** Inhibitory activities of LP-98 against HIV-1_BG505_ and HIV-1_JR-FL_ using a luciferase-based assay. **(D-F)** Analytical workflow for deriving binding kinetics between HIV-1 virions and LP-98 at different concentrations. Additional concentrations are shown in **Fig. S4**. **(D)** Representative single-molecule fluorescence intensity traces showing real-time binding between HIV-1 virions and LP-98 at 0.1, 1, and 5 nM. **(E)** Fitting of histograms of unbound dwell times yields observed rate constants (*K*_obs,i_). **(F)** Fitting of histograms of bound dwell times yields observed dissociation rate constants (*K*_off,i_). **(G)** Determination of the association rate constant (*K*_on_) from the linear dependence of *K*_obs,i_ on LP-98 concentration. The regression slope corresponds to *K*_on_. **(H)** Determination of the final dissociation rate constants (*K*_off_) as the mean value of *K*_off,i_ across all concentrations. **(I)** The equilibrium dissociation constant (binding affinity, *K*_d_) of LP-98 binding to HIV-1_BG505_-associated Env is 198 nM, calculated from the measured kinetic parameters.

Env on virions was labeled with a Cy5-derived fluorophore as described above, whereas LP-98 was labeled with a Cy3 at its N-terminus, leaving the C-terminal cholesterol group intact for membrane anchoring, a feature important for its antiviral potency^15,16,38^. Immobilized Cy5-labeled virions were incubated with Cy3-labeled LP-98 in the presence of host receptor soluble CD4 (sCD4) and coreceptor-mimicking antibody 17b to mimic the initial Env-host engagement environment that promotes Env trimer activation and helps release conformational constraints on gp41. We included sCD4 and 17b in all Env binding experiments. We tritiated virus-associated Env with LP-98 across concentrations from 0.1 to 5 nM. Representative fluorescence trajectories showed discrete association and dissociation binding events of LP-98 with virion-associated Env across all tested concentrations (**Fig. 2D** and **Fig. S4A**). Bound and unbound dwell times were extracted from individual trajectories and compiled into dwell-time histograms to determine off-rate, on-rate, and dissociation constant or binding affinity (*K*_off_, *K*_on_, and *K*_d_) (**Figs. 2E-I** and **Figs. S4B-C, S5**).

LP-98 exhibited rapid dissociation and high-nanomolar affinity for both HIV-1_BG505_ (**Fig. 2**) and HIV-1_JR-FL_ (**Fig. S5**). The measured (*K*_d_) values were ∼200 nM for HIV-1_BG505_ and 340 nM for HIV-1_JR-FL_. For both isolates, the rapid association and dissociation indicate dynamic, transient interactions of LP-98 with virion-associated Env. Of note, these high-nanomolar affinities contrast sharply with the sub-picomolar antiviral potency of LP-98, although they represent distinct biophysical and functional parameters and are not expected to correspond directly. This contrast suggests that prolonged Env occupancy (long residence time) is likely not required for fusion inhibition. The rapid kinetics are consistent with a “hit-and-run” mechanism, in which LP-98 could transiently engage Env and may repeatedly re-engage available Env targets. In the context of virus-to-cell fusion, the spatial constraints imposed by the close apposition of viral and cellular membranes may increase the effective local concentration of LP-98 that is conjugated with cholesterol for membrane anchoring and favor peptide rebinding. This may prolong the time LP-98 remains associated with the targeted Env beyond what our kinetic measurements imply. Such repeated encounters could also facilitate efficient interception of transient fusion-active Env conformations.

### LP-98 binding kinetics to native-like prefusion-stabilized Env trimers

For decades, it has been widely accepted that HR2-mimetic peptides bind to the putative gp41 prehairpin intermediate, in which HR1 is thought to form an exposed trimeric coiled coil resembling the HR1 core of the 6HB^2,21,39^. This model was inferred from inhibitor design principles and biochemical and structural studies of gp41 fragments^11,20,23,24^. However, whether these peptide inhibitors can engage the prefusion Env trimer has remained an open question. We therefore asked whether LP-98 can bind native-like prefusion Env SOSIP trimers. SOSIP is a widely used soluble form of Env ectodomain that introduces a disulfide bond between gp120 and gp41 (SOS) and the I559P substitution in gp41 (IP), facilitating the stabilization and purification of soluble prefusion Env trimers^40^. We purified the standard soluble BG505 SOSIP ectodomain, which is primarily in the prefusion closed conformation^41^ (Env_SS_; standard SOSIP; **Fig.S1**), and an A316W mutant SOSIP^42,43^(Env_MS_; mutant SOSIP; **Fig.S1**) by size-exclusion chromatography (SEC), followed by characterization using gel electrophoresis (SDS-PAGE), differential scanning fluorimetry (DSF), and negative stain electron microscopy (NSEM) (**Fig.3A** and **Fig.S6A**). Purified Env trimers were labeled according to established protocols^28–30^ (see **Methods**) to achieve predominantly single protomer dye incorporation, although a small fraction of trimers contained dyes on two or three protomers, as distinguished by stepwise photobleaching (**Fig. S7)**. We then measured LP-98 binding kinetics to both Env_SS_ and Env_MS_ trimers using our single-molecule approach (**Figs. 3B, 3C**, and **S6–S8**). Interestingly, LP-98 bound Env_ss_ with a *K*_d_ of 288 nM (**Figs. 3B** and **S8**), whereas stronger binding was observed for Env_MS_, yielding a *K*_d_ of 67 nM (**Figs. 3C** and **S6**). As with intact virions, rapid association and dissociation rates were observed for both soluble Env trimers, but Env_MS_ exhibited a longer residence time (**Figs. 3B, C**). As with virion-associated Env, LP-98 exhibited rapid association and dissociation kinetics when binding soluble Env trimers. These kinetics may reflect transient accessibility of the HR1-binding region within prefusion Env, in contrast to isolated gp41 HR1/NHR constructs that present a more readily accessible binding surface for HR2-derived peptides^11,24,26^. Together, these observations suggest that LP-98 binding is governed by conformational gating of HR1 accessibility within the trimer.

**Figure 3.**
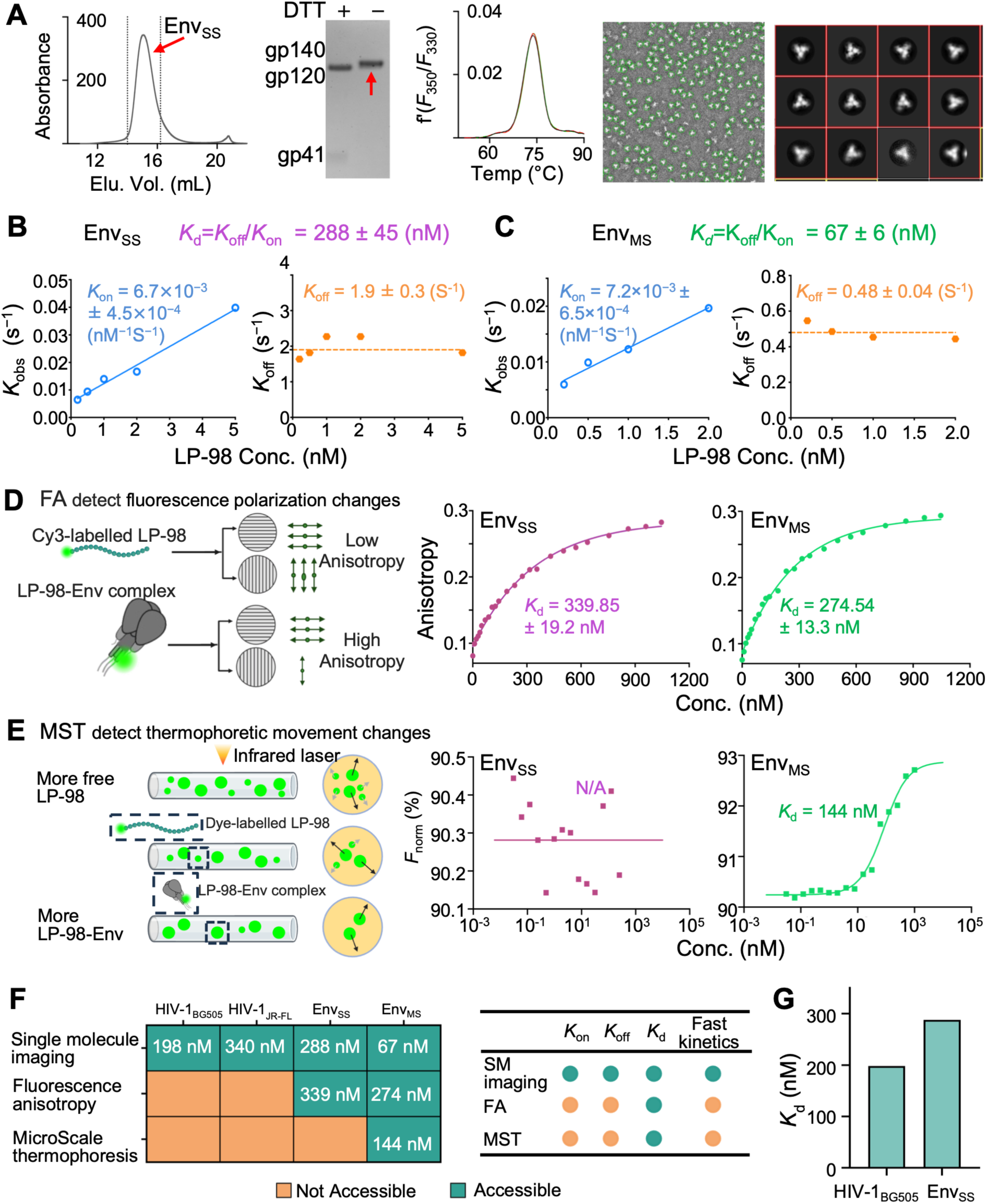
Single-molecule quantification of LP-98 binding kinetics to HIV-1 native-like prefusion Env trimers and comparison with bulk measurements. **(A)** Purification and characterization of prefusion-stabilized soluble Env trimers, Env_SS_, by size-exclusion chromatography (SEC), SDS–PAGE, differential scanning fluorimetry (DSF), and negative-stain electron microscopy (NSEM). **(B, C)** Single-molecule results of LP-98 binding kinetics to prefusion Env trimers: Env_SS_ (B) and Env_MS_ (**C**). **(D, E)** Binding affinity of LP-98 to Env_SS_ and Env_MS_ measured by FA (**D**) and MST (**E**). Fluorescence anisotropy (FA) measures changes in the rotational mobility of a fluorescently labeled molecule upon ligand binding, whereas microscale thermophoresis (MST) measures binding-induced changes in its thermophoretic movement within a temperature gradient. **(F)** Comparison of binding analysis methods. The single-molecule approach uniquely enables measurements of binding kinetics in the context of both intact virions and purified trimers, whereas FA and MST are limited to purified trimers and provide affinity measurements without kinetic information. **(G)** Affinity comparison of LP-98 binding to HIV-1_BG505_ and Env_SS_.

Using fluorescence anisotropy (FA; **Fig. 3D**), we measured a *K*_d_ of 339 nM for LP-98 binding to Env_SS_ and a *K*_d_ of 274 nM for binding to Env_MS_ (**Fig. 3D**), while microscale thermophoresis (MST; **Fig. 3E**) yielded a *K*_d_ of 144 nM for LP-98 binding to Env_MS_ but did not reliably detect binding to Env_SS_ under the tested conditions (**Fig. 3E**). These results are consistent with the stronger binding to Env_MS_ observed by single-molecule measurements (**Figs. 3B, 3C**). The unexpected binding of LP-98 to prefusion-stabilized soluble Env trimers expands the conformational window over which peptide inhibitors can engage Env beyond that predicted by the canonical prehairpin-intermediate model.

### Comparisons of LP-98 – Env binding across different contexts and approaches

Comparing LP-98–Env binding across different contexts and approaches revealed several notable findings. Our single-molecule approach enabled direct resolution of binding kinetics in the viral context that conventional bulk measurements cannot resolve (**Fig. 3F**). Comparison of virus-associated Env and soluble Env_SS_ revealed that LP-98 exhibits stronger binding to virus-associated Env than to the conformationally constrained soluble Env_SS_ trimer (**Fig. 3G**), which is stabilized to adopt prefusion conformations^41,44,45^. This finding is consistent with the notion that HR2-mimicking peptide inhibitors (such as LP-98) target the HR1 region of gp41^26^, which is largely buried in the prefusion closed state of Env and becomes progressively exposed during Env opening and subsequent gp41 refolding^21,22^.

Among the soluble Env trimers, Env_MS_, which carries the A316W substitution, showed stronger binding to LP-98 and slower dissociation than Env_SS_ (**Fig.3**). Previous studies^42,43^ have shown that A316W reduces V3 exposure and favors a prefusion closed Env trimer apex. At the same time, the SOSIP design constrains the downstream gp41 refolding transitions that would normally expose HR1, leaving the LP-98 binding region largely buried in the stabilized prefusion trimer. The increased LP-98 binding to Env_MS_, therefore, does not necessarily indicate a globally increased Env opening or full HR1 exposure. Instead, it may reflect altered conformational distributions, transient exposure of binding regions in prefusion intermediates, or asymmetric Env configurations^46–48^ in which individual protomers sample distinct states with differential binding accessibilities. To our knowledge, the binding kinetics and affinity of LP-98 with gp41 fragments, soluble Env trimers, or virus-associated Env have not previously been reported. Our results provide the first such kinetic characterization. In contrast, prior studies of HR2-mimicking fusion inhibitors binding to isolated HR1/NHR or engineered gp41 targets such as 5-Helix have reported a broad range of affinities, including *K*_d_ values of ∼30 nM for T20 to 5-Helix^23^, ∼200 nM – low μm for C34 to N36 and 5-Helix^49,50^, and ∼0.0007 nM for C37 to 5-Helix^23^, compared with ∼60 - 400 nM for LP-98 measured here across different trimeric Env contexts and approaches (**Figs. 2 - 3**). These differences likely reflect, at least in part, how HR1 is presented and accessed, which can differ substantially between isolated or engineered targets and intact Env.

### T-118 and 4ca binding kinetics to RSV native-like prefusion-stabilized F trimers

To assess whether our findings and platform extend beyond HIV-1 Env, we next examined peptide inhibitor binding to the RSV fusion (F) protein (**Fig. 4A**), another class I viral fusion protein that similarly mediates entry through large-scale conformational rearrangements. Analogous to the HIV-1 Env HR1 region, the HRA region of RSV F serves as the target for HRB-derived peptide fusion inhibitors (**Fig. 4A**). Peptides such as T-118 and 4ca inhibit viral entry by disrupting 6HB formation during F refolding (**Figs. 4A-B**) and exhibit potent antiviral activities^18,19^. We tested T-118 using a fluorescence plaque assay and revealed a low-nanomolar IC_50_ (**Figs. 4C and S9**). Nevertheless, neither T-118 nor 4ca binding kinetics to F have been characterized.

**Figure 4.**
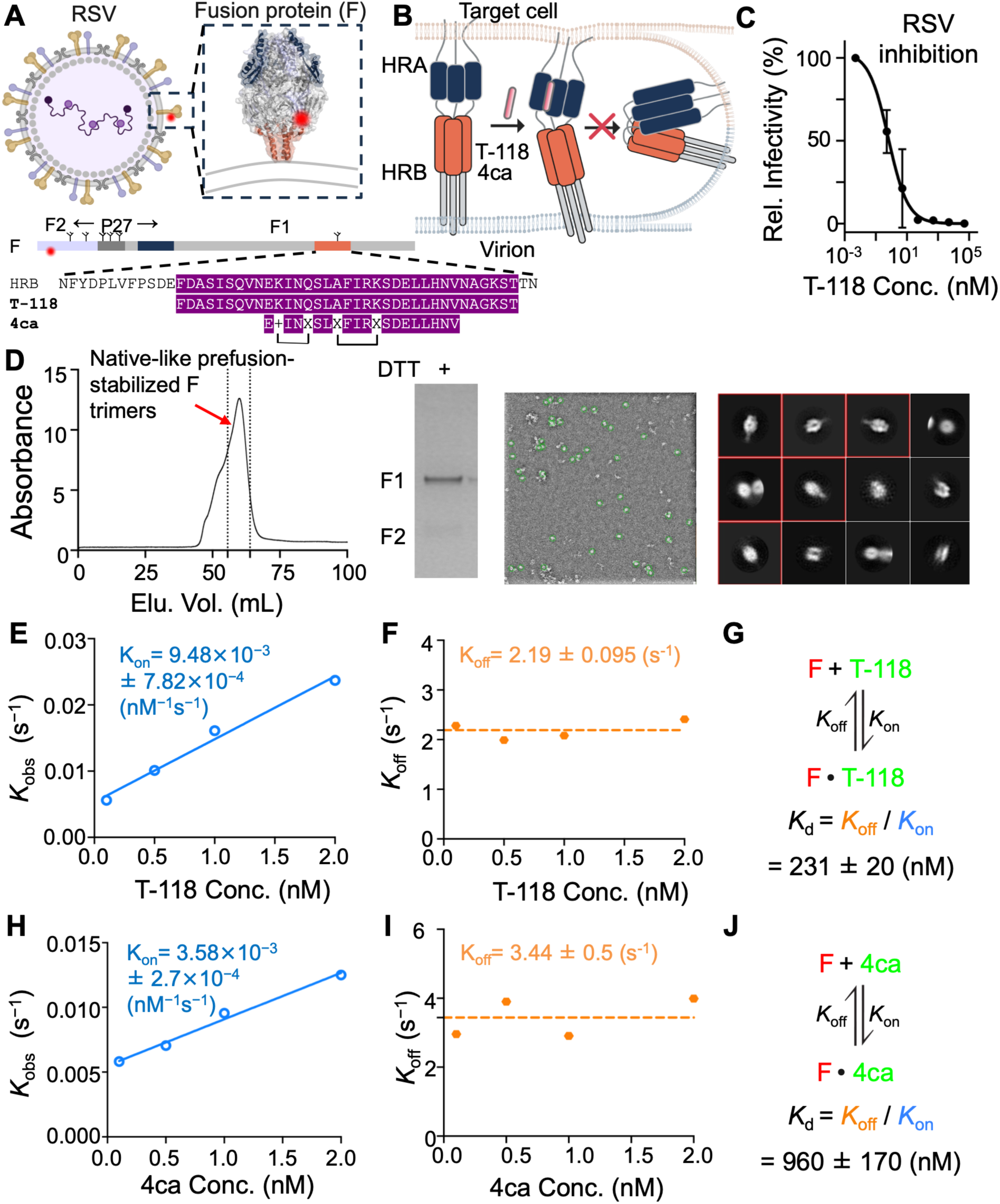
Single-molecule quantification of T-118 binding kinetics to RSV native-like prefusion F trimers. **(A)** Structure of a Cy5-labeled RSV fusion (F) glycoprotein trimer in its prefusion conformation (PDB ID: 4JHW). F is composed of F1 and F2 subunits. The heptad repeat A (HRA) region is the target of HRB-derived fusion-inhibitory peptides T-118 and 4ca, with conserved residues highlighted. **(B)** The widely accepted mechanism of RSV fusion inhibition by T-118 and 4ca. HRB-derived peptide inhibitors are thought to target a putative prehairpin intermediate, as proposed for class I viral fusion proteins. **(C)** Inhibitory activities of T-118 against RSV quantified from infected Hep2 cells using a fluorescence plaque assay (**Fig.S9**). **(D)** Purification and characterization of prefusion-stabilized soluble F trimer ectodomains by SEC, SDS–PAGE, and NSEM. **(E - G)** Binding kinetics of T-118 to prefusion F trimers characterized using single-molecule imaging, including *K*_on_ (**D**), *K*_off_ (**E**), and *K*_d_ (**F**). **(H - J)** Binding kinetics of 4ca to prefusion F trimers characterized using single-molecule imaging, including *K*_on_ (**G**), *K*_off_ (**H**), and *K*_d_ (**I**).

We purified a native-like prefusion-stabilized F trimer (**Fig. S3**)^51^ and characterized its morphology by SEC, SDS-PAGE, and NSEM (**Fig. 4D**). This prefusion F trimer is the major target of neutralizing antibodies; as such, current FDA-approved RSV vaccines Arexvy (GSK)^52,53^ and Abrysvo (Pfizer)^54,55^ are based on this prefusion F design^51,56,57^. To apply our single-molecule approach, we introduced a short peptide tag into F for Cy5-derived dye labeling (**Fig. S3**). We then directly quantified interactions between Cy3-labeled T-118 or 4ca and prefusion F trimer by measuring binding across a range of peptide concentrations (**Figs. S10** and **S11**). Analysis of bound and unbound dwell times yielded a *K*_d_ of 231 nM for T-118 binding to prefusion F (**Figs. 4E – G)** and 960 nM for 4ca (**Figs. 4H – J**). Both T-118 and 4ca exhibited rapid association and dissociation kinetics (**Figs. 4E – J**), similar to what we observed for LP-98 binding to HIV-1 Env.

Together, these findings show that peptide fusion inhibitors targeting two distinct class I viral fusion systems, HIV-1 Env and RSV F, engage prefusion-stabilized trimers with rapid and reversible kinetics in the high-nanomolar affinity range. Consistent with our observations for HIV-1 Env, these results further suggest that peptide inhibitor engagement is not restricted to the canonical prehairpin intermediate and can occur while fusion proteins remain in prefusion-like conformations.

### Predominance of 1:1 peptide-to-trimer binding stoichiometry

Because our approach monitors real-time binding to individual soluble Env or F trimers, it allows us to resolve peptide-to-trimer stoichiometry directly from the fold increase in Cy3 fluorescence intensity of individual binding bursts. In principle, binding of two ligands (2:1) to one trimer generates fluorescence bursts with approximately twice the intensity of single-ligand binding events (1:1), whereas binding of three ligands (3:1) gives rise to bursts with approximately threefold greater intensity (**Fig. 5A**).

**Figure 5.**
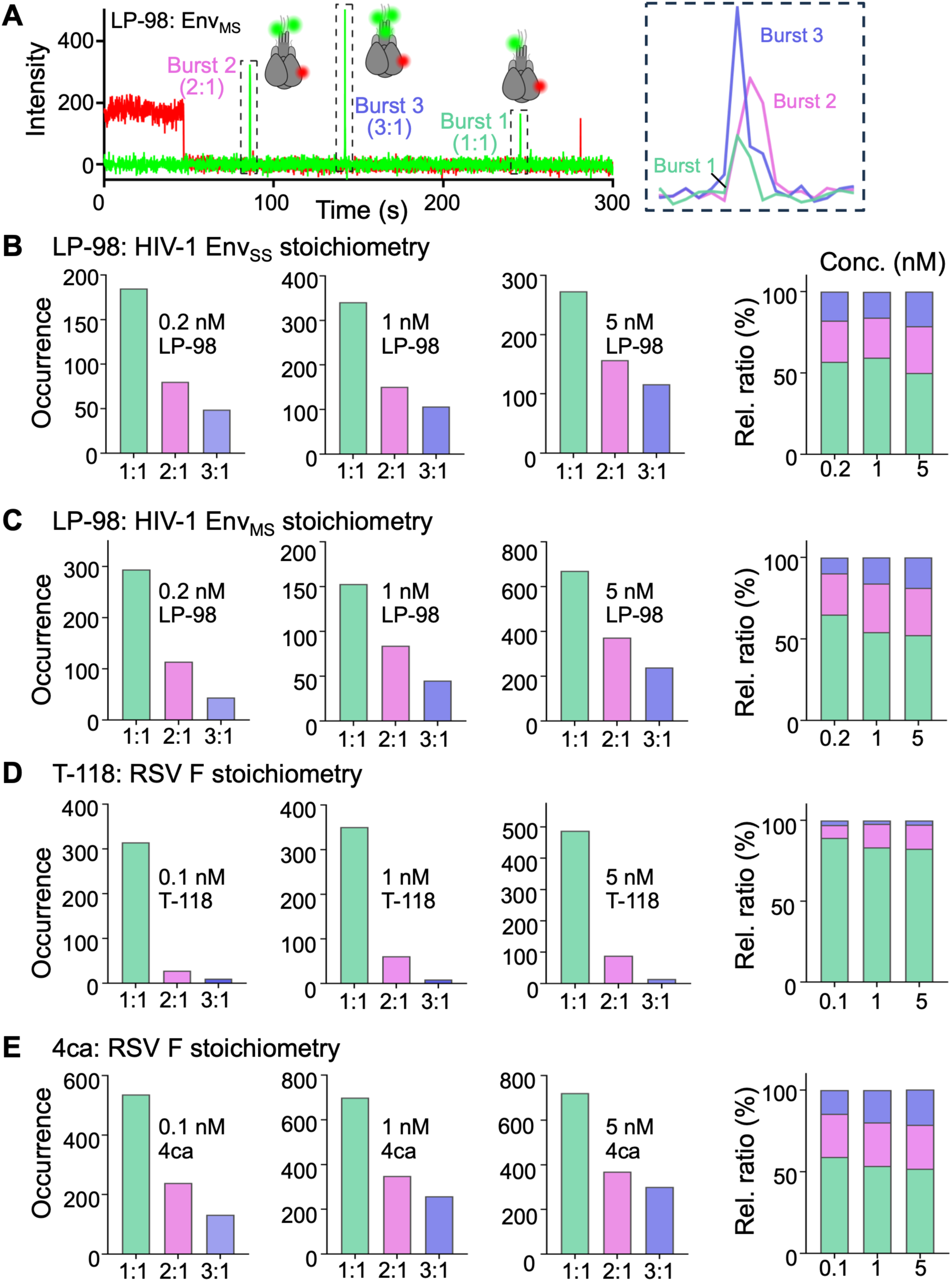
A predominant 1:1 peptide-to-trimer binding stoichiometry emerges from heterogeneous occupancy distributions. **(A)** Representative fluorescence trajectory showing distinct binding occupancies of Cy3-labeled LP-98 on a Cy5-labeled Env_MS_. Three binding events (dashed boxes) exhibit discrete fluorescence intensity levels corresponding to the binding of one, two, or three LP-98, as illustrated in the accompanying schematics. **(B-E)** Binding stoichiometry of LP-98 binding to Env_SS_ (**B**) and Env_MS_ (**C**), and of T-118 (**D**) and 4ca (**E**) binding to F. Binding events corresponding to one-peptide (1:1, green), two-peptide (2:1, magenta), and three-peptide (3:1, blue) occupancy were quantified at each peptide concentration. The far-right panel summarizes the normalized fractions of each occupancy state across peptide concentrations.

Using this approach, we analyzed fluorescence bursts arising from LP-98 binding to Env_SS_ and Env_MS_ across low, intermediate, and high concentrations and quantified binding stoichiometry. We observed all three possible peptide-to-trimer stoichiometries (1:1, 2:1, and 3:1) for both Env trimers (**Figs. 5B, C**). Notably, 1:1 peptide-to-trimer stoichiometry predominated in the heterogeneous population (>50%), whereas 2:1 and 3:1 stoichiometries together accounted for less than or close to 50% of observations (**Figs. 5B, C**). The predominance of 1:1 stoichiometry persisted across peptide concentrations, suggesting that it is largely independent of ligand concentration.

We next applied the same analysis to T-118 and 4ca binding to RSV F trimers. Consistent with the LP-98 results, 1:1 stoichiometry again represents the predominant population in heterogeneous distributions (**Figs. 5D, E**). This preference is even more striking for T-118, with less than 20% of events corresponding to 2:1 or 3:1 stoichiometries in total (**Fig. 5D**). For 4ca, the distributions of stoichiometry ratio across different peptide concentrations are very similar to those of LP-98 to Env trimers (**Fig.5E**). These observations suggest that full trimer occupancy is not the dominant mode of peptide fusion inhibitor engagement, at least for systems tested here.

Thus, despite the threefold symmetry of HIV-1 Env and RSV F trimers and the potential for binding of up to three peptide molecules, peptide engagement was characterized by heterogeneous occupancy distributions in which single-peptide occupancy predominated. This pattern was observed across peptide concentrations and in both viral fusion systems examined here, indicating that full trimer occupancy is not the dominant mode of peptide engagement under the conditions tested.

### Multiple routes of sequential and simultaneous binding lead to higher peptide occupancy

We next examined how the less frequent 2:1 and 3:1 peptide-to-trimer stoichiometries observed in our experiments arise. Using our single-molecule imaging approach, we directly visualized individual association (on) and dissociation (off) events, allowing us to distinguish simultaneous and sequential/stepwise binding modes in real time.

We observed four possible routes (or binding modes) to and from higher-peptide occupancy states, representing different combinations of simultaneous and stepwise association and dissociation: simultaneous association/simultaneous dissociation, stepwise association/simultaneous dissociation, simultaneous association/stepwise dissociation, and stepwise association/stepwise dissociation. Representative fluorescence trajectories from LP-98 binding to soluble Env trimers illustrate these four modes. The most frequently observed route was simultaneous association followed by simultaneous dissociation (**Fig. 6A**). Additional trajectories revealed stepwise association followed by simultaneous dissociation (**Fig. 6B**), simultaneous association followed by stepwise dissociation (**Fig. 6C**), and stepwise association followed by stepwise dissociation (**Fig. 6D**).

**Figure 6.**
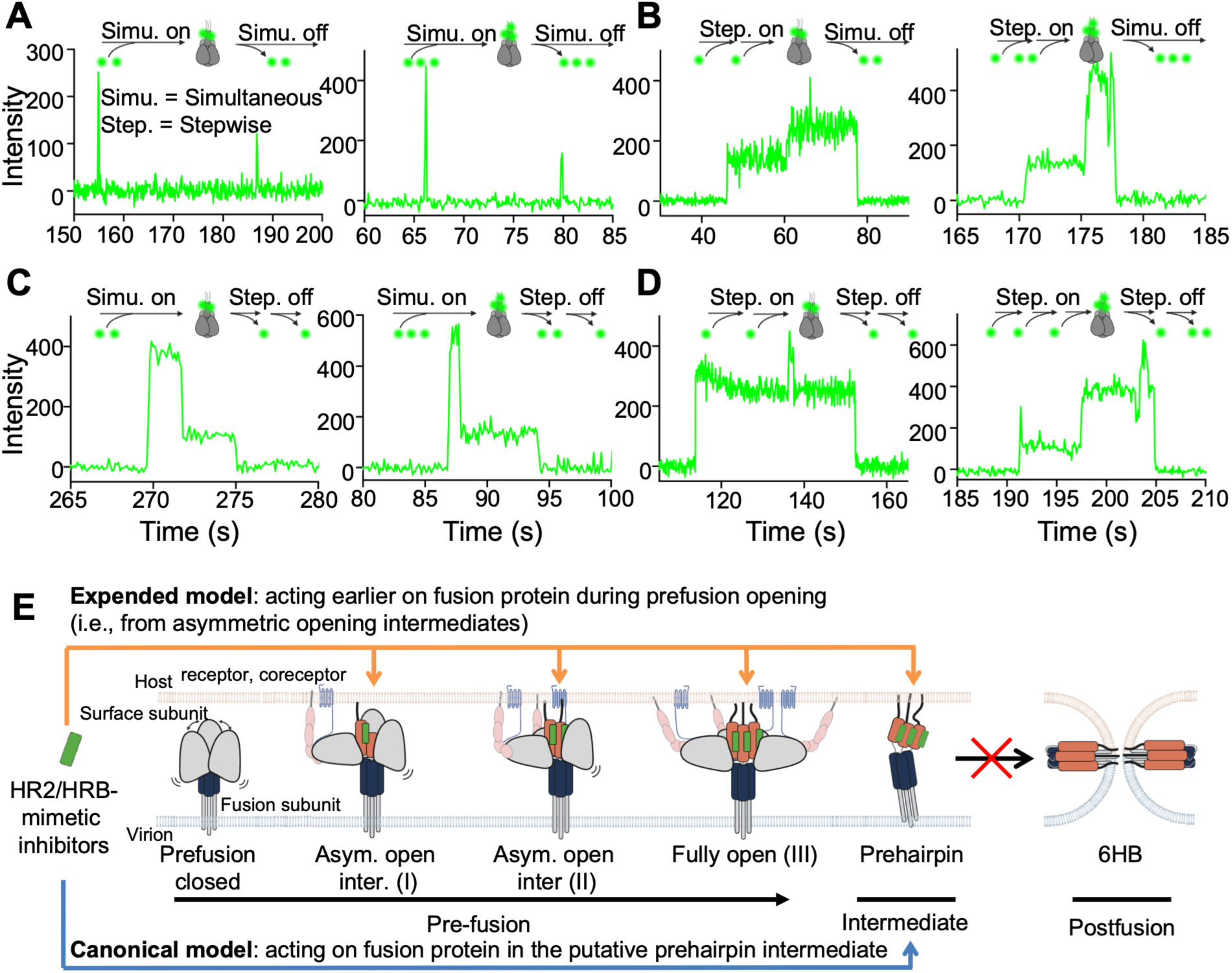
Multiple binding routes to form high peptide-trimer occupancy and the derived working model from this study. (**A**) Representative trajectories showing simultaneous association (on) and simultaneous dissociation (off) of two inhibitors (left) or three (right) binding to a single fusion protein trimer. Corresponding binding modes are illustrated in the accompanying schematics. Examples are from LP-98 and the Env trimer. (**B-D**) Representative trajectories, as in panel A, showing the other three possible binding modes for two (left panel) or three LP-98 (right panel) binding to a single trimer, including stepwise association with simultaneous dissociation (**B**), simultaneous association with stepwise dissociation (**C**), and stepwise association with stepwise dissociation (**D**). (**E**) HR2/HRB-mimetic peptide inhibitors act on fusion proteins earlier during prefusion than the canonical putative prehairpin intermediate, illustrated with virus-associated Env and receptor/coreceptor presenting host membranes.

Similar binding behaviors were observed across all peptide-trimer systems examined. Thus, higher-occupancy states can form and decay through multiple association and dissociation routes, revealing substantial heterogeneity in peptide fusion inhibitor engagement. Whether these binding events involve cooperativity between protomers within individual trimers remains to be determined.

## Discussion

In this study, we used single-molecule imaging to quantify the binding kinetics and stoichiometry of HR2/HRB-derived peptide fusion inhibitors interacting with class I viral fusion proteins. Our findings reveal dynamic and heterogeneous binding behaviors across two viral systems, HIV-1 Env and RSV F. Specifically, we directly resolved peptide inhibitor binding kinetics on virions, identified unexpected engagement of prefusion viral fusion proteins, observed a persistent predominance of single-peptide occupancy despite the threefold symmetry of Env and F trimers, and uncovered multiple routes involving sequential and simultaneous binding events.

In addition to revealing previously unrecognized features of peptide engagement, our single-molecule approach enabled direct quantification of inhibitor binding kinetics to native Env displayed on intact virions, a measurement that has been difficult to achieve using conventional ensemble methods. By preserving Env in its native membrane and viral context, this approach provides kinetic information that more closely reflects interactions at the virion surface. This is evident in our observation that LP-98 exhibits stronger apparent binding to virus-associated Env than to conformationally constrained, prefusion-stabilized soluble Env trimers.

Direct observation of peptide inhibitor binding to prefusion Env and F trimers was unexpected. The widely accepted model of heptad-repeat-derived peptide fusion inhibition proposes that these inhibitors act primarily after activation of the prefusion trimer, when the N-terminal heptad-repeat helical region becomes exposed in the transient prehairpin intermediate and accessible to inhibitors^10,11,21–23^. For HIV-1 Env, HR2/CHR-derived peptides bind the exposed HR1/NHR region of gp41, preventing association with the endogenous HR2/CHR and formation of the stable 6HB^10,11,20,23,24,26^. A similar model has been proposed for RSV F or other class I fusion proteins, in which HRB-derived peptides engage the HRA region during F refolding and interfere with 6HB formation^18,19^. Our observation that peptide inhibitors bind prefusion-stabilized Env and F trimers indicates that their binding can begin earlier than this canonical model proposes (**Fig. 6E**). Such prefusion engagement does not necessarily imply global opening of the fusion protein or complete exposure of the HR1/HRA region. Increased LP-98 binding to Env_MS_ may therefore reflect the conformational dynamics of prefusion Env, in which local HR1 accessibility and protomer asymmetry create opportunities for peptide engagement without requiring global Env opening. Such asymmetric or transient accessibility may permit peptide binding to prefusion trimers.

This finding does not exclude the prehairpin intermediate as an important inhibitory target, but instead broadens the fusion-stage window over which peptide inhibitors may first target viral fusion proteins (**Fig. 6E**). In line with this, previous studies have raised the possibility that HIV-1 HR2-derived inhibitors can engage gp41 beyond a putative prehairpin intermediate, including others progressing toward 6HB formation^58^. This possibility is also consistent with 6HB-like structures formed by gp41 fragments in complex with HR2-derived peptides^11,20,24–26^. Whereas previous studies suggested that peptide inhibitors may remain engaged beyond a putative prehairpin intermediate and into later stages of refolding, our results extend the accessible window in the opposite direction by revealing interactions with prefusion-stabilized trimers. Thus, rather than acting exclusively on a fusion intermediate, peptide fusion inhibitors can begin engaging viral fusion proteins earlier in the fusion pathway than previously appreciated (**Fig. 6E**). Such prefusion interactions may have remained unresolved because peptide inhibitors were designed to bind complementary helical regions of the fusion machinery and were therefore commonly characterized using isolated fusion-protein fragments, while historical challenges in solubilizing and stabilizing viral fusion protein trimers further limited studies in the prefusion trimeric context.

A second unexpected finding is the predominance of single-peptide occupancy in peptide-to-trimer stoichiometry despite the threefold symmetry of Env and F trimers. The trimeric architecture of class I viral fusion proteins and the canonical 6HB model permits binding of three peptide fusion inhibitor molecules per trimer, as shown in structural studies^11,24–26^. In contrast, our single-molecule measurements reveal heterogeneous peptide occupancy of soluble prefusion trimers, with single-peptide occupancy predominating over two- and three-peptide occupancy. This pattern persists across peptide concentrations and is observed for inhibitors targeting both HIV-1 Env and RSV F, suggesting that it is not simply a consequence of low ligand availability or a feature unique to a single fusion protein. Thus, peptide occupancy of prefusion trimers does not simply follow the threefold symmetry expected from the 6HB structure. It may arise from conformational asymmetry, differential accessibility among protomers constrained in prefusion, positive or negative cooperativity, or kinetic constraints on subsequent binding, although the present data do not distinguish among these possibilities. These factors may act together to shape the observed occupancy distributions. The sequential and simultaneous binding events observed at higher occupancies further indicate that peptide engagement is kinetically heterogeneous and can proceed through multiple pathways to reach similar occupancy states.

Overall, our findings reveal an expanded targetable fusion-stage window and previously inaccessible binding mechanisms of peptide fusion inhibitors, broadening the canonical model of class I viral fusion inhibition and potentially informing the development of improved inhibitors. Key technical limitations include the inability to resolve binding stoichiometry on intact virions and the finite kinetic range accessible by single-molecule imaging. Despite these limitations, our approach enables direct characterization of peptide inhibitor binding to both virus-associated Env and soluble Env and F trimers at the level of individual events. Future studies integrating single-molecule binding with structural and functional measurements could define the molecular basis of heterogeneous occupancy, test for binding cooperativity, and establish how these binding behaviors relate to antiviral inhibition.

Intrigued by key findings in this work, our observations suggest a more dynamic model of peptide fusion inhibition than traditionally envisioned. Rather than engaging fusion proteins only after formation of a discrete prehairpin intermediate, peptide inhibitors may target multiple conformational states along the fusion pathway, including prefusion-stabilized trimers and later fusion intermediates. In this framework, inhibition may arise from repeated transient encounters with accessible target regions rather than prolonged occupancy of a single conformational state.

## Methods and Materials

### Cell lines and cell maintenance

Human embryonic kidney 293T (HEK293T) cells (ATCC, CRL-3216) were used to produce replication-defective HIV-1 viral particles. TZM-bl cells (BEI Resources, HRP-8129) were used as target cells for HIV-1 infection, and Hep-2 cells (ATCC, CCL-23) were used for RSV infection. Cells were maintained in high-glucose Dulbecco’s Modified Eagle Medium (DMEM; Gibco, Cat. No. 11965-092) supplemented with 10% (v/v) fetal bovine serum (Gemini Bio, Cat. No. 100-106), 100 U/mL penicillin, 100 μg/mL streptomycin (Gibco, Cat. No. 15140-122), and 2 mM L-glutamine (Gibco, Cat. No. 25030-081). Cells were cultured at 37°C in a humidified incubator with 5% CO_2_.

FreeStyle 293-F cells (Thermo Fisher Scientific, Cat. No. R79007) were used for transient expression of HIV-1 Env trimer, and cells were maintained in FreeStyle 293 Expression medium (Thermo Fisher Scientific, Cat. No. 12338018). Expi293F cells (Thermo Fisher Scientific, Cat. No. A39240) were used for transient expression of RSV F trimer, 17b, and soluble CD4 molecules, and cells were maintained in Expi293F Expression Medium (Gibco, Cat. No. A1435102) according to the manufacturer’s instructions. Cells were cultured in suspension at 37°C in a humidified atmosphere containing 8% CO_2_ with continuous orbital shaking.

### Plasmid construction

The Q23_BG505_N136* plasmid used to produce HIV-1_BG505_ viral particles was generated by site-directed mutagenesis using Amber-free HIV-1_Q23_ Env_BG505_ ΔRT (Addgene #213007) as a template^28^. In this construct, ΔRT denotes deletion of the gene encoding reverse transcriptase (RT)^28,46,59^. The Amber-free construct contains TAA stop codons in place of all preexisting TAG (amber) stop codons in the Pol, Vif, Vpu, and Rev genes, providing a background suitable for site-specific amber suppression in HIV-1, including BG505 Env^28^. To generate Q23_BG505_N136*, the codon corresponding to N136 in Env was replaced with a TAG stop codon. The resulting plasmid was amplified in Stbl3 competent cells (Invitrogen, #C7373-03) and verified by DNA sequencing.

The Env-encoding plasmid (JRFL_N136*) used to produce HIV-1_JR-FL_ viral particles was created from the pCAGGS vector and wild-type JRFL Env as a template, introducing the N136* substitution as previously described^60,61^. We used the packaging plasmid pCMV-dR8.2 (Addgene, Cat. No. 12263) along with the Env-encoding plasmid in a 1:1 ratio during transfection to produce virions suitable for single-molecule imaging.

Plasmids used to produce soluble Env trimer were generated using BG505 SOSIP.664 gp140 as a template (Env_SS_), with the addition of a polyhistidine tag at the C-terminus for purification^40^. We introduced the A316W substitution into Env to generate Env_MS_. The peptide A4 (DSLDMLEW) was introduced into Env for dye labeling (**Fig.S1**). The soluble F trimers were constructed based on a his-tagged plasmid RSV-F DS-Cav1^56^ (a generous gift from Peter D. Kwong’s laboratory), and RSV-F DS-Cav1-Q3 was generated by substituting T36 with the Q3 peptide (GQQQLG) for F labeling (**Fig. S3**).

### Production and fluorescent labeling of HIV-1 virions

The preparation of HIV-1_BG505_ and HIV-1_JR-FL_ viral particles has been previously described^27,46,62^. 24 h before transfection, healthy, exponentially growing, mycoplasma-free HEK293T cells were seeded into culture plates to reach>70% confluency at the time of transfection. The growth medium was then replaced with Opti-MEM (Gibco, Cat. No. 31985-070), and plasmid DNA was mixed with polyethylenimine (PEI, 1 mg/mL) at a 3:1 PEI: DNA ratio (v:w). The mixtures were incubated for 15 min at room temperature before adding to cells. An additional plasmid that encodes an amber stop codon suppressor, tRNA^pyl^/NESPyIRS^AF^ (a gift from the Edward Lemke Lab), was co-transfected to enable incorporation of the noncanonical amino acid (ncAA^63,64^). This plasmid encodes the orthogonal tRNA/synthetase pair that incorporates ncAA into its cognate tRNA and was co-transfected at a 3:1 ratio relative to the Env plasmid. The ncAA trans-cyclooct-2-en-L-lysine (TCO*A; SiChem, Cat. No. SC8008) was added to the culture medium at a final concentration of 250 μM. After 4 - 6 h of incubation, the transfection medium was replaced with growth medium, and cells were incubated for an additional 40 - 48 hours. Supernatants containing viral particles were collected, filtered through a 0.45 μm membrane, and concentrated by ultracentrifugation through a 15% (w:v) sucrose in PBS at 25,000 rpm for 2 h using an SW28 rotor (Beckman Coulter).

Viral particle pellets were resuspended in labeling buffer (50 mM HEPES pH 7.0, 10 mM CaCl_2_, 10 mM MgCl_2_) and incubated overnight at room temperature with 0.2 μM tetrazine-conjugated Cy5 derivative (LD655-TTZ, Lumidyne) fluorophore^28,29^. The reaction was quenched by adding 1 μM BCN-OH followed by incubation for 10 minutes. DSPE-PEG2000-biotin (Avanti Research, Cat. No. A88129) was then added to a final concentration of 0.1 mg/mL, and the mixture was incubated for 30 minutes at room temperature with rotation. Excess dye was removed by ultracentrifugation on a 6–18% OptiPrep gradient for 1 hour at 40,000 rpm at 4 °C using a SW40Ti rotor (Beckman Coulter). Gradient fractions containing labeled virions were collected, aliquoted, and stored at −80 °C until use in imaging experiments.

### Production of recombinant soluble HIV-1 Env and RSV F trimers

The preparation of fluorescently labeled Env through the enzymatic labeling peptide tag (A4, DSLDMLEW) has been described previously^27^. The A4 tag is tolerated at specific sites in the V4 variable region of gp120 without affecting Env expression, processing, virus incorporation, or virus infectivity, as previously identified^46^. Env proteins were produced by transient transfection of suspension-adapted FreeStyle 293-F cells with a mix of ectodomain-encoding tag-free and a trace amount of A4-tagged Env plasmids. Prior to transfection, cells were maintained in expression medium and diluted to a density of 1.5 × 10⁶ cells/mL. Plasmid DNA encoding Env was mixed with furin protease at a 4:1 mass ratio, followed by mixing with PEI diluted in Opti-MEM. The mixture was incubated at room temperature for 20 min before adding to cells. Transfected cells were then cultured at 37 °C with 8% CO₂ under agitation at 120 rpm for 5 days. Culture media were centrifuged at 4,000 rcf for 30 min, and the supernatant was filtered through 0.22 μm polyethersulfone (PES) membranes before loading onto a PGT145 IgG-conjugated affinity column that was preequilibrated in 20 mM PBS, pH 7.5. Following extensive washing, bound Env trimers were eluted using 3 M MgCl₂, pH 7.2. Eluted proteins were concentrated to approximately 1 mL using a centrifugal filter (Amicon® Ultra Centrifugal Filter, 100 kDa). Samples were subsequently filtered through 0.22 μm PES membranes to remove aggregates prior to size-exclusion chromatography on a Superose 6 Increase 10/300 GL column (Cytiva) pre-equilibrated in PBS, using an AKTA Pure system. Fractions corresponding to trimeric Env were pooled, concentrated, snap frozen in liquid nitrogen, and stored at −80 °C.

For F production, a plasmid encoding the F ectodomain was co-transfected with furin at a 4:1 ratio into suspension-adapted Expi293F cells, followed by 5-day culture in expression medium as described above. Culture media were collected by centrifugation at 4,000 × g for 30 min, and the supernatant was filtered through 0.22 μm PES membranes before loading to a Ni²⁺-nitrilotriacetic acid (Ni-NTA) affinity chromatography. Bound F was washed and eluted using buffer containing 20 mM Tris-HCl (pH 8.0), 200 mM NaCl, and 250 mM imidazole. Further purification was performed by size-exclusion chromatography (SEC) using HiLoad 16/600 Superdex 200 column pre-equilibrated with 2 mM Tris-HCl (pH 7.5), 150 mM NaCl, and 0.02% sodium azide. Fractions corresponding to RSV F trimers were pooled and concentrated, snap-frozen in liquid nitrogen, and stored at −80 °C.

Protein samples were analyzed by SDS–PAGE under standard conditions and visualized by Coomassie Brilliant Blue staining. Protein thermal stability was assessed by differential scanning fluorimetry (DSF), and melting temperatures were determined from the corresponding thermal unfolding profiles.

### Fluorescent labeling of recombinant soluble Env and F trimers

Dye labeling of soluble Env or F trimers was performed, as previously described for virus-associated Env^46,62^, purified Env trimers^27^, or SARS-CoV-2 spike proteins^65^. Env carrying a genetically encoded A4 tag was enzymatically labeled by AcpS (acyl carrier protein synthase)^66^ with the Cy5 derivative fluorophores (LD655-Coenzyme A; LD655-CoA, Lumidyne Technologies), whereas F carrying a genetically encoded Q3 tag was enzymatically labeled by transglutaminase (TGase)^67^ with the Cy5 derivative fluorophores (LD655-cadaverine; LD655-CD, Lumidyne Technologies) in the labeling buffer described above. Labeling reactions were performed at room temperature for 6 h. Excess free dyes were removed using Zeba dye and biotin removal columns (Thermo Fisher Scientific, Cat. No. A44299). For immobilization on passivated, streptavidin-coated microfluidic imaging chambers, Env and F were incubated with biotinylated anti-His antibody (BIO-RAD, Cat. No. MCA1396B) at 4°C overnight.

### Production of soluble CD4 and 17b

Soluble CD4 (sCD4) and 17b were transiently expressed and purified as described previously^68^. In brief, sCD4 was transiently expressed in Expi293F cells and purified using Q425-affinity chromatography, followed by size-exclusion chromatography on a HiLoad 16/600 Superdex 200 pg column equilibrated in 20 mM PBS, pH 7.5, 0.002% w/v sodium azide. 17b was expressed in Expi293F cells by transient transfection of heavy-light chain plasmids and purified with protein A affinity chromatography and size-exclusion chromatography using a HiLoad 16/200 Superdex 200 pg column equilibrated in 20 mM PBS, pH 7.5, 0.002% w/v sodium azide. Purified sCD4 and 17b were buffer-exchanged to 20 mM PBS, pH 7.5, 0.002% w/v sodium azide, then snap-frozen for use.

### Negative staining electron microscopy (NSEM)

A frozen aliquot of sample stored at −80 °C was thawed at room temperature for 5 min and diluted to 30 µg/mL in HBS (20 mM HEPES, 150 mM NaCl, pH 7.4) supplemented with 0.02 g/dL ruthenium red and 8 mM glutaraldehyde. After 5 min of incubation, the crosslinking reaction was quenched by adding 1 M Tris (pH 7.4) to a final concentration of 80 mM. The samples were then applied to glow-discharged carbon-coated grids for 8-10 s, blotted, and washed twice with 1/20× HBS. Grids were stained with 2 g/dL uranyl formate for 1 min, blotted, and air-dried. Data were acquired on a Philips EM420 microscope at 120 kV and 49,000x magnification. Forty micrographs were recorded using a 76-megapixel CCD camera at a pixel size of 2.4 Å. Image processing and 2D classification were performed in Relion 3.0.

### Microscale Thermophoresis (MST)

The Env complex was prepared by incubating Env with 17b Fab and sCD4 in a 1:5 molar ratio for 60 hours at room temperature in PBS buffer, pH 7.4. LP-98 was biotinylated using the Biotin Labeling Kit (Cat. No. NT-L020, Biotinylated Target Labeling Kit, NanoTemper Technologies) according to the manufacturer’s instructions. For MST assay, biotinylated LP-98 was used at a fixed concentration of 20 nM and titrated against a two-fold serial dilution series of the Env complex. A total of 16 concentrations of Env complex, ranging from 1 µM to 30.5 pM, were prepared in MST buffer containing 50 mM Tris (pH 8.0), 80 mM NaCl, 0.05% Tween-20, and 0.3 mM TCEP. The mixtures were incubated for an additional 24 hours at room temperature before loading into Monolith NT.115 capillaries. Binding interactions between LP-98 and the Env complex were monitored and measured using the Monolith NT.115 instrument (NanoTemper Technologies). Binding affinity data were analyzed using NanoTemper Affinity Analysis software (version 2.3).

### Fluorescence anisotropy (FA)

100 nM of Cy3-labeled LP-98 (100 nM) peptide was incubated with increasing concentrations of Env (10 nM to 1200 nM) at room temperature for 30 min before data acquisition. Fluorescence anisotropy measurements were acquired using a Horiba FluoroMax spectrofluorometer (FLUOROMAX_PLUS_C) with appropriate excitation and emission settings for Cy3. Parallel and perpendicular fluorescence intensities were recorded, and anisotropy values were calculated using the instrument software. Binding curves were generated by plotting anisotropy values against Env concentration and fitted using a one-site binding model to determine the equilibrium dissociation constant.

### HIV-1 inhibition assay

The antiviral activity of LP-98 against HIV-1 was determined using a luciferase-based viral inhibition assay. Env-pseudotyped HIV-1 was generated using the envelope glycoprotein construct encoding Env_BG505_ or Env_JRFL_ and a packaging construct encoding backbone pNL4-3.Luc.R-E-(BEI Resources, Cat # HRP-3418,)^69,70^, which carries a firefly luciferase reporter. TZM-bl cells were seeded in 96-well plates at 2 × 10^4^ cells per well 24 hours before the assay. LP-98 was serially diluted in 10-fold increments with four replicates. The diluted peptide was mixed with HIV-1 virus (10^5^ RLU) and incubated at 37 °C for 1 hour. The peptide-virus mixture was then transferred onto the pre-seeded TZM-bl cells. Following a 48-hour incubation, luciferase activity was measured using a BioTek Synergy H1 multimode microplate reader. The dose-response curve was fitted with a four-parameter variable-slope logistic regression model in GraphPad.

### RSV inhibition assay

The antiviral activity of T-118 against RSV was determined using a fluorescence plaque assay. The rgRSV224 virus (Cat # NR-52018, BEI Resources)^71^, an RSV A2 strain carrying a GFP reporter, was used for the assay. Hep-2 cells were seeded in 96-well plates at 2 × 10^4^ cells per well 24 hours before the assay. T-118 was serially diluted in 10-fold increments, with three per dilution. The peptide was mixed with rgRSV224 at a multiplicity of infection (MOI) of 0.01 and incubated at 37 °C for 1 hour. The mixture was then transferred to the pre-seeded Hep-2 cells. Following a 48-hour incubation, the GFP-positive area in each well was quantified using a BioTek Cytation C10 confocal imaging reader. The dose-response curve was fitted using a four-parameter variable-slope logistic regression model in GraphPad.

### Single-molecule ligand-binding data acquisition

All single-molecule ligand-binding image stacks were acquired on an in-house-built prism-based total internal reflection fluorescence microscope. The reaction chamber was customized and passivated with a mixture of PEG and biotin–PEG, which was coated with streptavidin (Invitrogen). LD655 (Cy5-derived dye)-labeled virions or fusion proteins were then immobilized on passivated streptavidin-coated stop-flow imaging chambers, followed by application of a Cy3-labeled peptide. The data acquisition steps are essentially the same as previously described for smFRET studies^27,28,30,62^, except that these experiments were specifically designed to avoid FRET; therefore, we used excitation from two single-wavelength lasers instead of one. In ligand-binding, Cy3 and LD655 fluorophores were simultaneously excited by the evanescent field generated by total internal reflection of a 532-nm continuous-wave (CW) diode-pumped solid-state laser (Ventus, Laser Quantum) and a 633-nm CW solid-state laser (Obis, Coherent). The two fluorescence signals were collected through a water-immersion 60× 1.27-NA objective (Nikon), spectrally split by a MultiCam LS image splitter (Cairn Research) equipped with a dichroic filter (Chroma) and directed through ET590/50 and ET690/50 emission filters (Chroma) corresponding to Cy3 and Cy5 channels, respectively. Image stacks or movies were recorded using synchronized sCMOS cameras (Hamamatsu ORCA-Flash 4.0 V3) at 10 frames per second for 300 seconds, using custom software implemented in LabView (National Instruments). All data acquisitions were performed in a buffer containing 50 mM Tris (pH 7.4), 50 mM NaCl, a cocktail of triplet-state quenchers and 2 mM protocatechuic acid (PCA) with 8 nM protocatechuic-3,4-dioxygenase (PCD) to remove molecular oxygen^72,73^. Where indicated, Env was incubated with 0.1 mg/mL of sCD4 and 0.1 mg/mL of 17b for 30 min at room temperature prior to addition of LP-98. After a further 30 min incubation with the peptide, data were acquired. The procedure was repeated with increasing concentrations of LP-98 (0.1–5 nM). For RSV F experiments, except for sCD4 and 17b, all other procedures were identical.

### Single-molecule ligand-binding data analysis

Fluorescence trajectories were extracted from collected image stacks or movies using the customized MATLAB (Mathworks) program SPARTAN^74^. Trajectories were manually inspected, and only those containing signals from both channels, with Cy5 fluorescence intensities ≥100 units above baseline and at least one binding event indicated by a Cy3 signal, were selected for further analysis. The selected trajectories were analyzed in Igor (WaveMetrics), where bound and unbound dwell times were defined and measured.

For each ligand concentration (*i*), dwell times (*t*) were compiled into a histogram, and the resulting distribution was fitted in Igor using a single-exponential decay model to obtain the time constant (*τ_i_*).

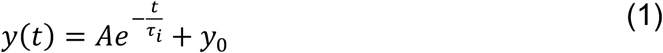

where *A* is the amplitude and *y*_0_ is the offset.

In this manuscript, dissociation time constant (*τ*_off,i_) is reported as *τ*_off,i_ ± SE_τo*FF*,i_, where SE_τo*FF*,i_ represents the standard error of the fitted time constant obtained from the nonlinear regression.

Dissociation rate constant at each ligand concentration (*K*_off,i_) were then calculated as:

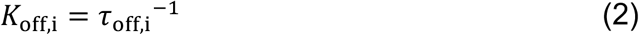

In this manuscript, *K*_off,i_ is presented as *K*_off,i_ ± SE_off,i_, where SE_off,i_ represents propagated standard error derived from the uncertainty of the fitted dissociation time constant.

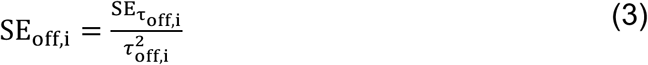

As *K*_off_ is independent of ligand concentration, the final *K*_off_ value was reported as the mean of the values obtained across all tested ligand concentrations (calculated and plotted using GraphPad Prism):

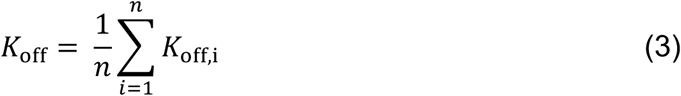

In this manuscript, *K*_off_ is reported as *K*_off_ ± SEM_off_, where SEM_off_ represents the standard error of the mean of *K*_off,i_ across the tested ligand concentrations.

Similarly, for each ligand concentration (*i*), unbound dwell times (*t*) were compiled into a histogram and fitted with a single-exponential decay to derive association time constant (*τ*_on,i_) using Eq. (1) in Igor. In this manuscript, association time constant (*τ*_on,i_) is reported as *τ*_on,i_ ± SE_τon,i_, where SE_τon,i_ represents the standard error of the fitted time constant obtained from the nonlinear regression. The observed association rate constant (*K*_obs,i_) was calculated as:

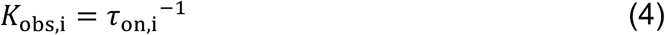

In this manuscript, *K*_obs,i_ is reported as *K*_obs,i_ ± SE_obs,i_, where SE_obs,i_ represents the propagated standard error derived from the uncertainty of the fitted association time constant.

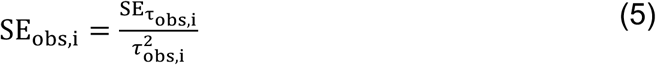

*K*_obs_ reflects the frequency of subsequent binding events and is proportional to ligand concentration. Based on this relationship, the association rate constant ( *K*_on_) was determined as (calculated and plotted using GraphPad Prism):

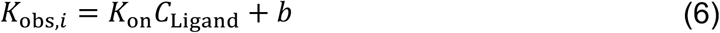

where b represents the fitted y-intercept and *C*_Ligand_ denotes the ligand concentration. In this manuscript, *K*_on_ is reported as *K*_on_ ± SE, where SE represents the standard error of the slope obtained from the linear regression (calculated using GraphPad Prism).

The equilibrium dissociation constant (*K*_d_) was calculated from *K*_on_ and *K*_off_ as:

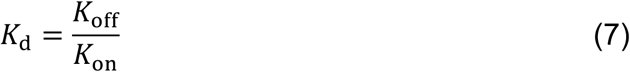

In this manuscript, the equilibrium dissociation constant is reported as 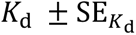, where 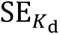 represents the standard error of *K*_d_ and is calculated using SEM_off_ and SE according to the error propagation equation.

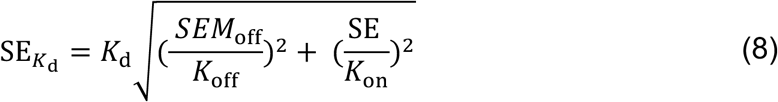

## Data Availability

Data supporting the findings of this study are available within the paper and its supplementary information files.

## Acknowledgements

The following reagents were obtained through BEI Resources, NIAID, NIH: Recombinant Respiratory Syncytial Virus, A2 Expressing Green Fluorescent Protein (GFP) (rgRSV224), NR-52018; Human Immunodeficiency Virus Type 1 (HIV-1) NL4-3 ΔEnv Vpr Luciferase Reporter Vector (pNL4-3.Luc.R-E-), HRP-3418. We thank Peter D. Kwong for generously providing the tag-free RSV F construct used in this study, and thank Edward Lemke for generously providing the amber suppressor construct. The HIV-1-related studies were supported by the National Institute of Allergy and Infectious Diseases of the National Institutes of Health (NIH) under Award Number R01 AI181600 (M.L.) The RSV-related studies were supported by the National Institute of General Medical Sciences of the National Institutes of Health (NIH) under Award Number R35 GM151169 (M.L.). Purification of HIV-1 Env ectodomains, NSEM and MST analysis were supported by the National Institute of Allergy and Infectious Diseases of the National Institutes of Health (NIH) under Award Number U54 AI170752 (P.A.). The content is solely the responsibility of the authors and does not necessarily represent the official views of the NIH.

## Author contributions

M.L. conceived the study. M.L. and P.A. supervised the study. M.L., R.H.K., and J.L. designed the experiments. Y.H., R.H.K., and J. L. performed single-molecule binding experiments. R.H.K., Y.H., and J.L. analyzed single-molecule binding data. R.H.K. and Y.H. performed fluorescence anisotropy measurements. W.X. performed virus inhibition experiments. R.H.K. and N.K.G. performed mutagenesis, prepared viral particles, and performed site-specific dye labeling. R.H.K., X.H., and B.T. performed protein purification. R.J.E. performed NSEM analysis. J. J. L. and B.T. performed MST measurements. K.J. and Y.H. assisted with cell maintenance, plasmid preparation, and study coordination. J.L. and R.H.K. prepared the figures. M.L., J.Y., and R.H.K. wrote the manuscript with input from all authors.

---

**Figure S1.**
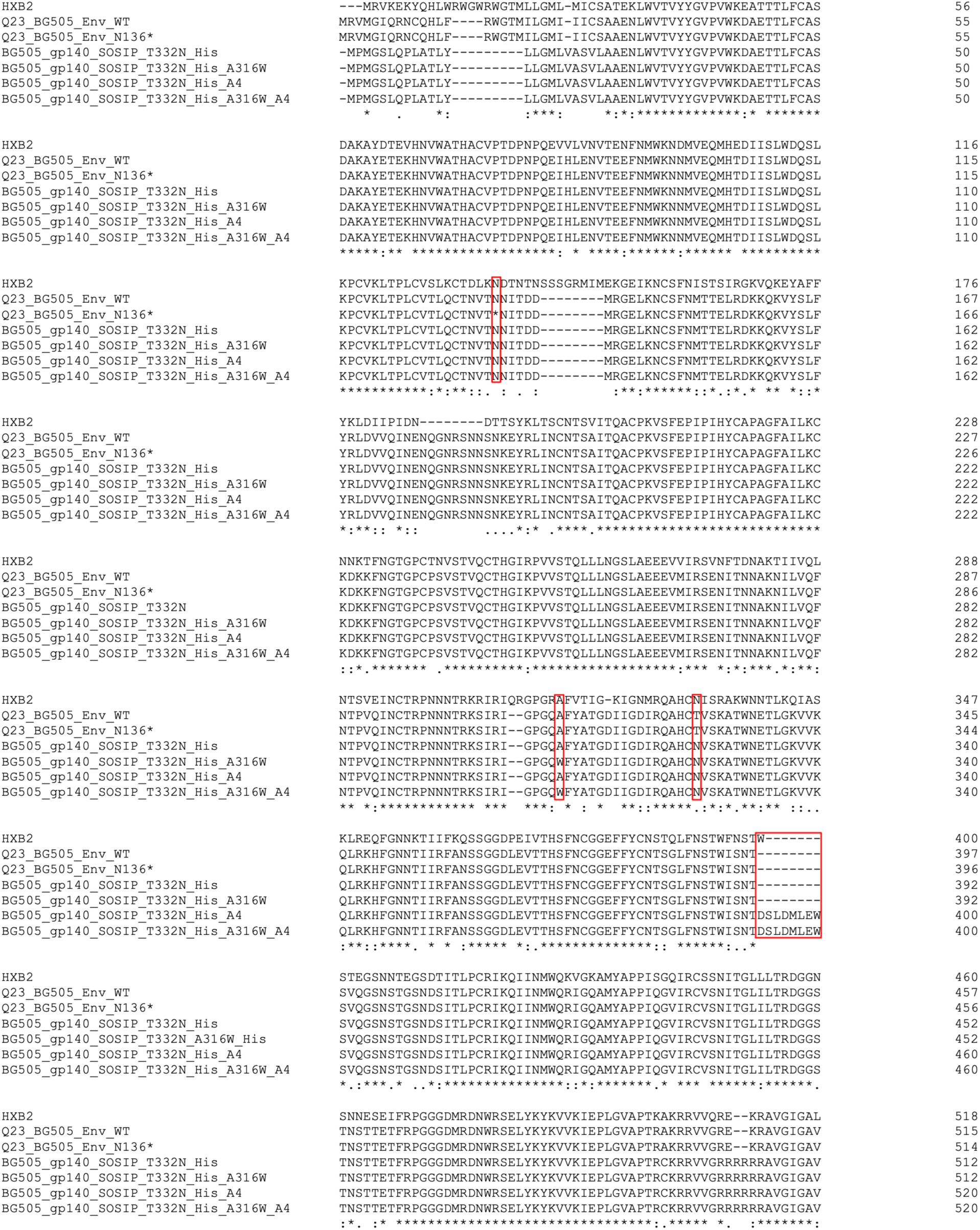
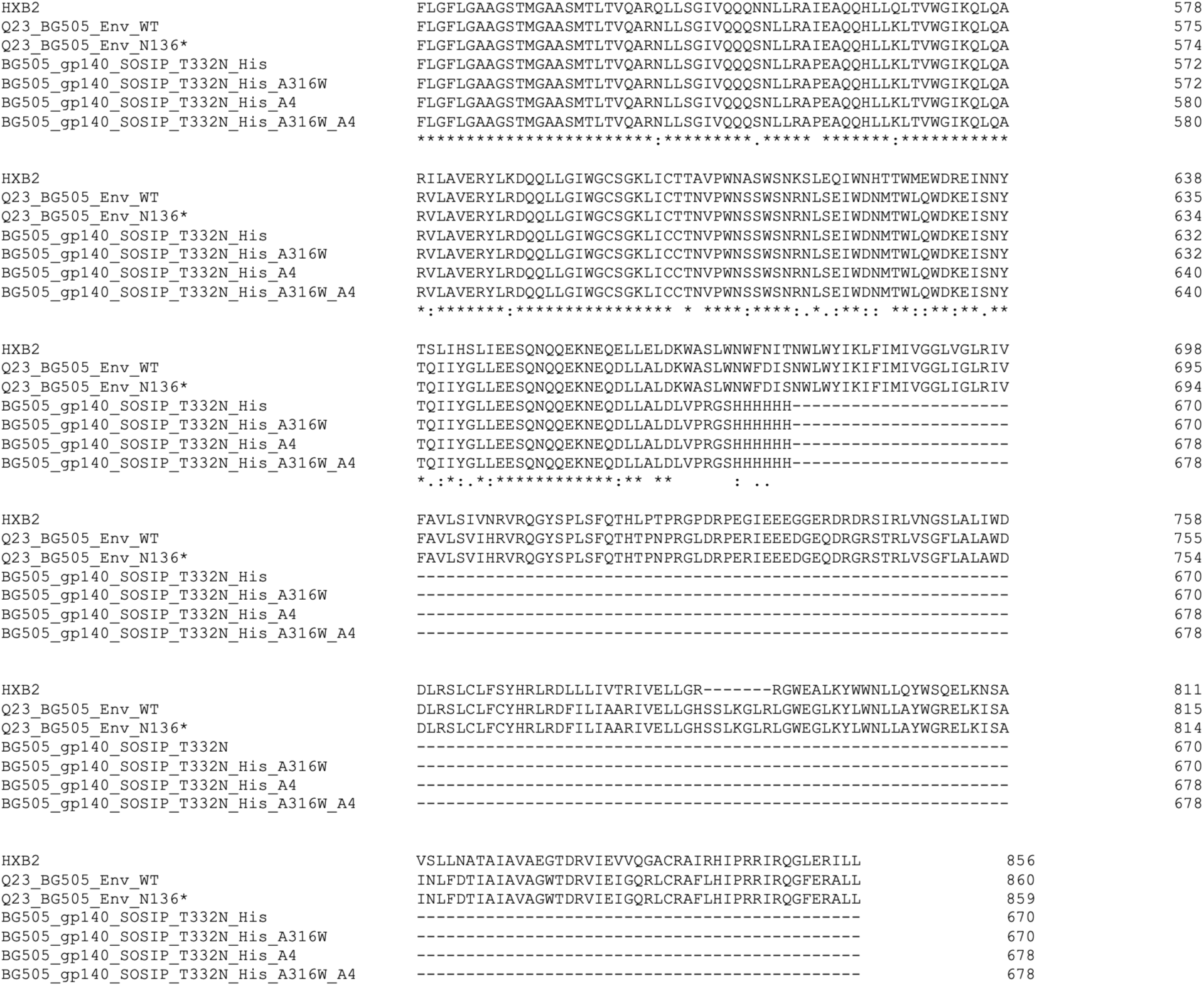
Sequence alignment of BG505 Env constructs used in this study. Q23_BG505_Env indicates virus-associated BG505 Env HIV-1_BG505_, with the suffix “WT” indicating wild type and “N136*” indicating an amber codon at position 136 for click-dye labeling with an ncAA. BG505_gp140_SOSIP_T332N_His denotes soluble BG505 SOSIP ectodomains (Env_SS_), and BG505_gp140_SOSIP_T332N_His_A316W denotes the variant carrying the additional A316W substitution (Env_MS_). The suffix “A4” denotes the peptide tag. The Cy5-labeling sites at N136* and A4, and the substitution A316W, are highlighted in red boxes. Residues are numbered based on the HXB2 reference sequence.

**Figure S2.**
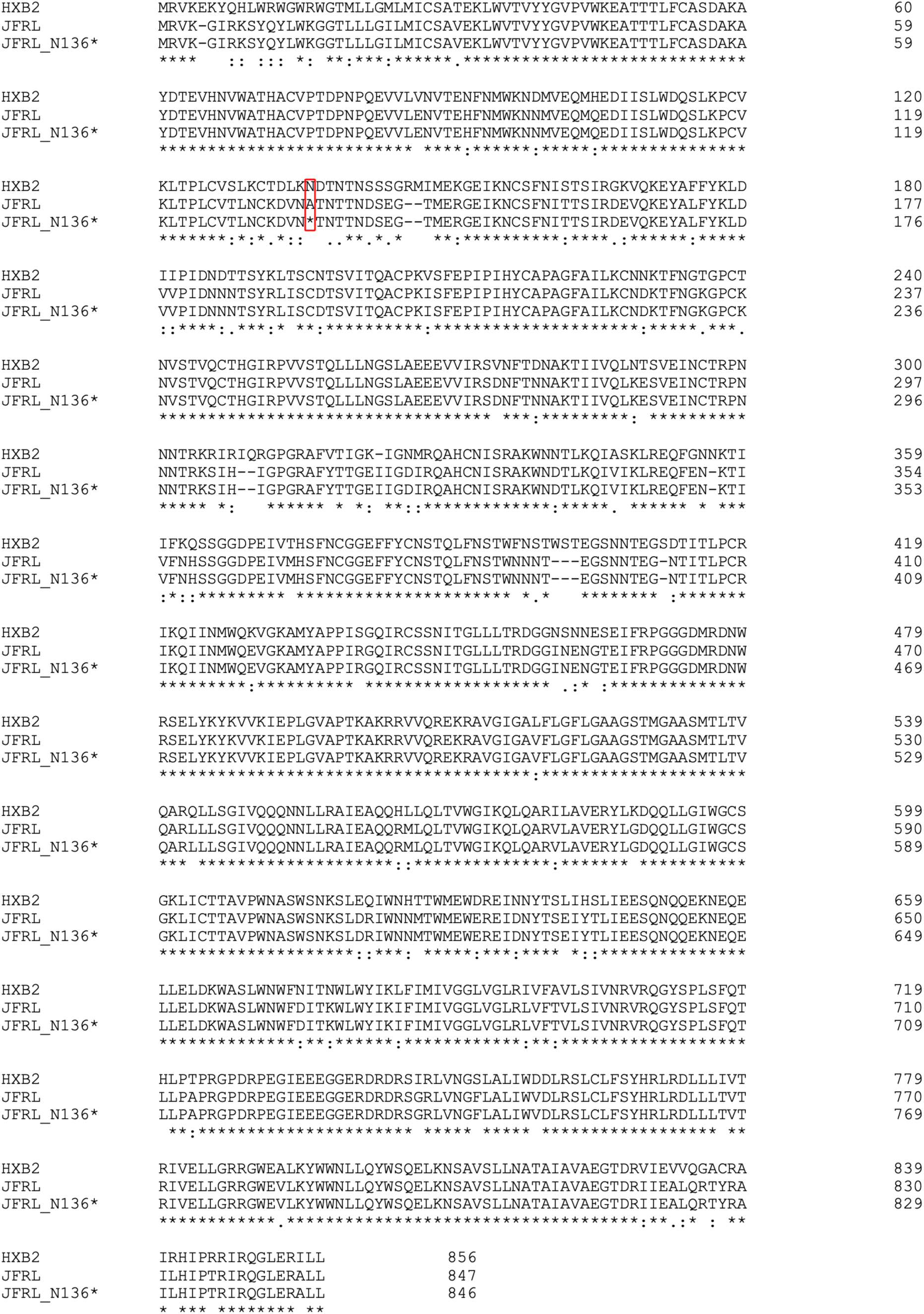
Sequence alignment of tag-free and tagged JR-FL Env. Virus-associated JR-FL Env was used in this study. JRFL denotes the wild-type Env, whereas N136* indicates the introduced amber codon for click dye labeling with an ncAA. The Cy5-labeling site at N136* is highlighted. Residues are numbered based on the HXB2 reference sequence.

**Figure S3.**
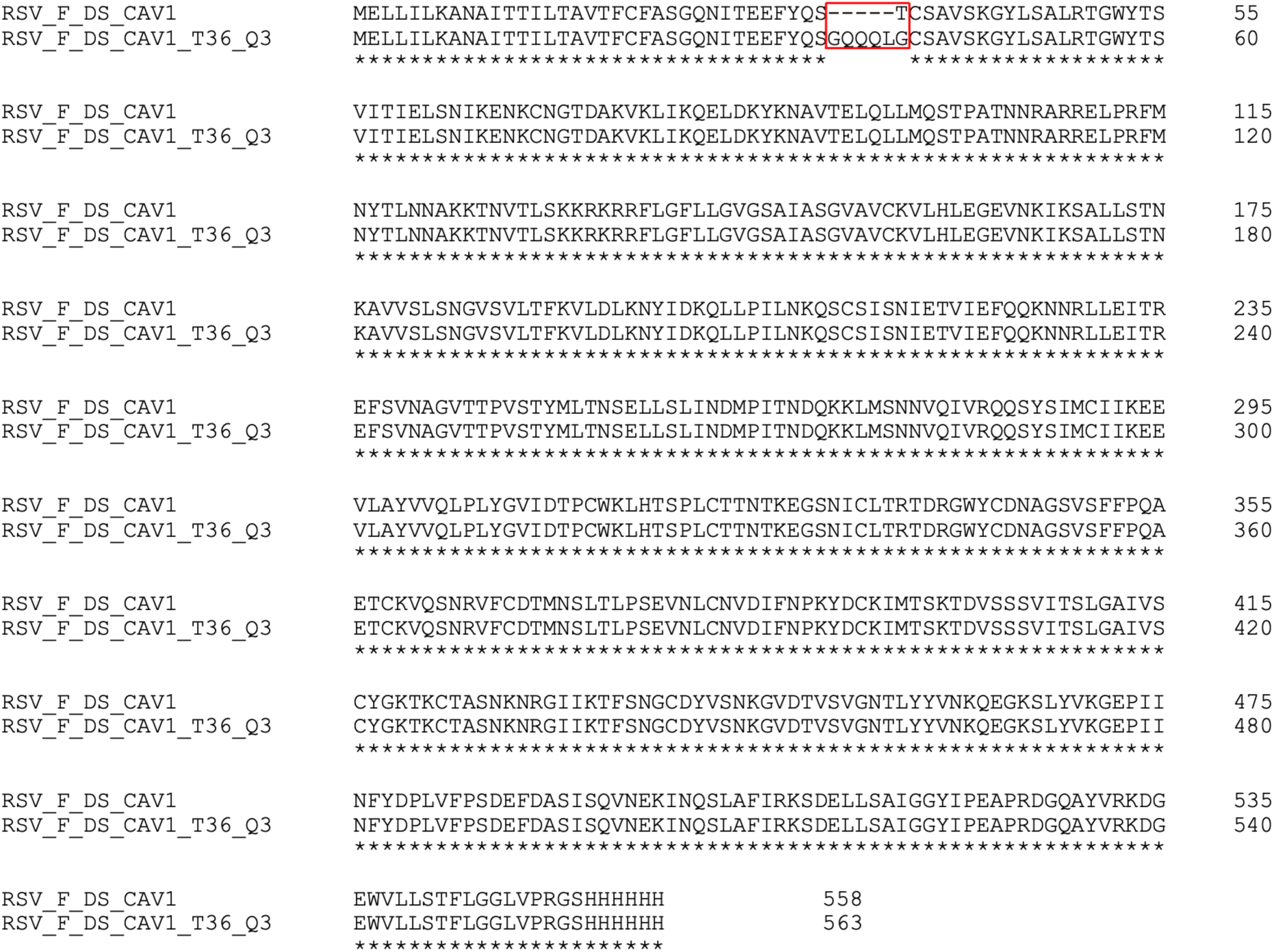
Sequence alignment of soluble RSV F ectodomains. RSV_F_DS_CAV1 represents the parental F, and RSV_F_DS_CAV1_T36_Q3 refers to the Q3-tagged version for fluorescence labeling. The Cy5-labeling site (T36_Q3) is highlighted.

**Figure S4.**
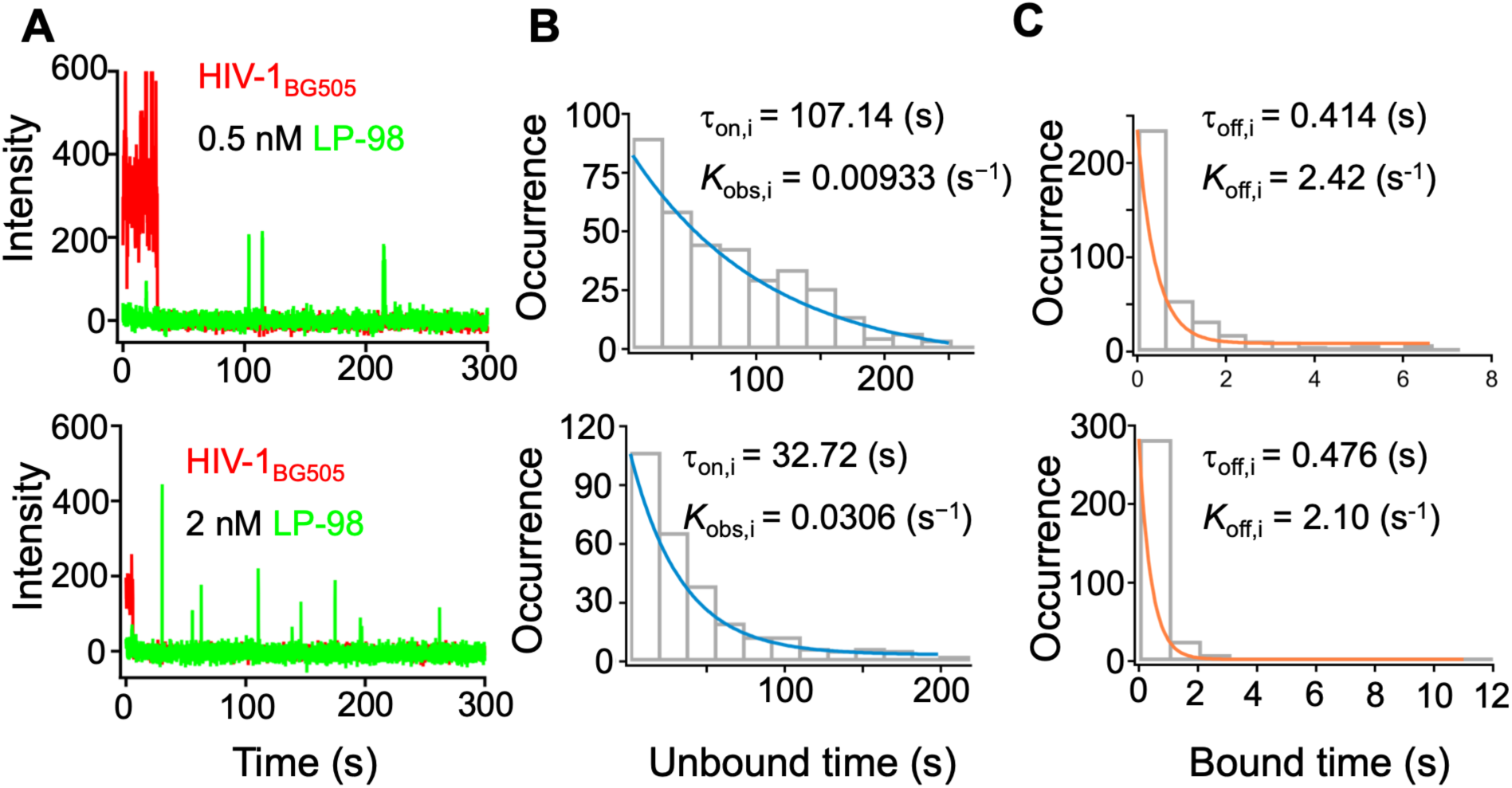
Additional peptide concentrations used for single-molecule measurements of LP-98 binding kinetics to HIV-1_BG505_ (others are shown in Fig. 2). **(A-C)** Results as in Figs. 2D-E, respectively, for measurements performed at two additional peptide concentrations included in the kinetic titration series. Top panels: 0.5 nM LP-98. Bottom panels: 2 nM LP-98.

**Figure S5.**
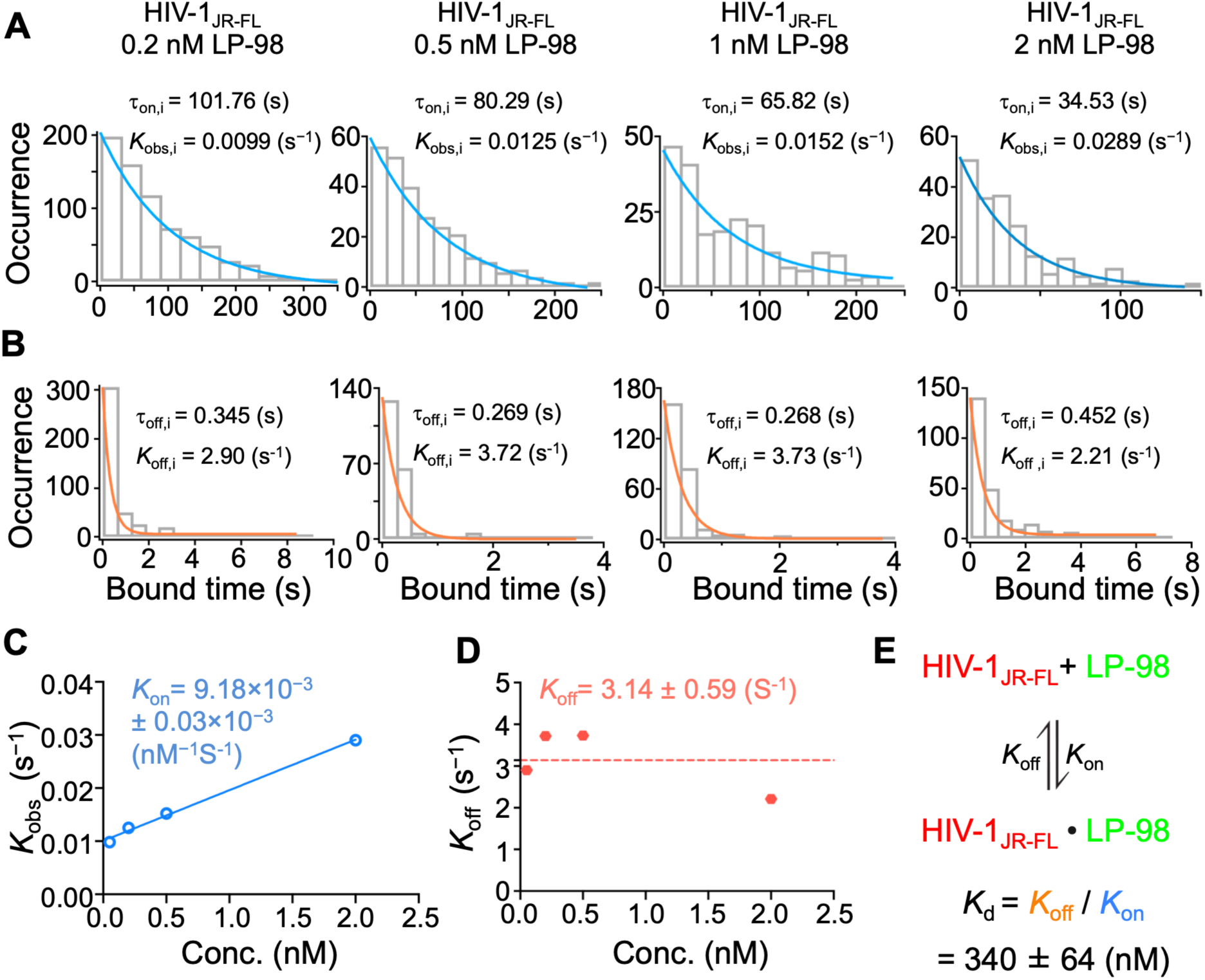
Single-molecule quantification of LP-98 binding kinetics to HIV-1_JR-FL_. **(A)** Histograms of unbound dwell times fitted with single-exponential decays yield *K*_obs,i_ values that increase with LP-98 concentration (0.2nM, 0.5nM, 1nM, and 2nM). **(B)** Histograms of bound dwell times yield *K*_off,i_ values that are independent of LP-98 concentration, consistent with first-order dissociation. **(C)** *K*_obs,i_ plotted as a function of LP-98 concentration; linear fitting yields *K*_on_. **(D)** Determination of the final dissociation rate constants (*K*_off_) as the mean value of *K*_off,i_ across all tested LP-98 concentrations. **(E)** Estimated *K*_d_ for LP-98 binding to HIV-1_JR-FL_ virion-associated Env.

**Figure S6.**
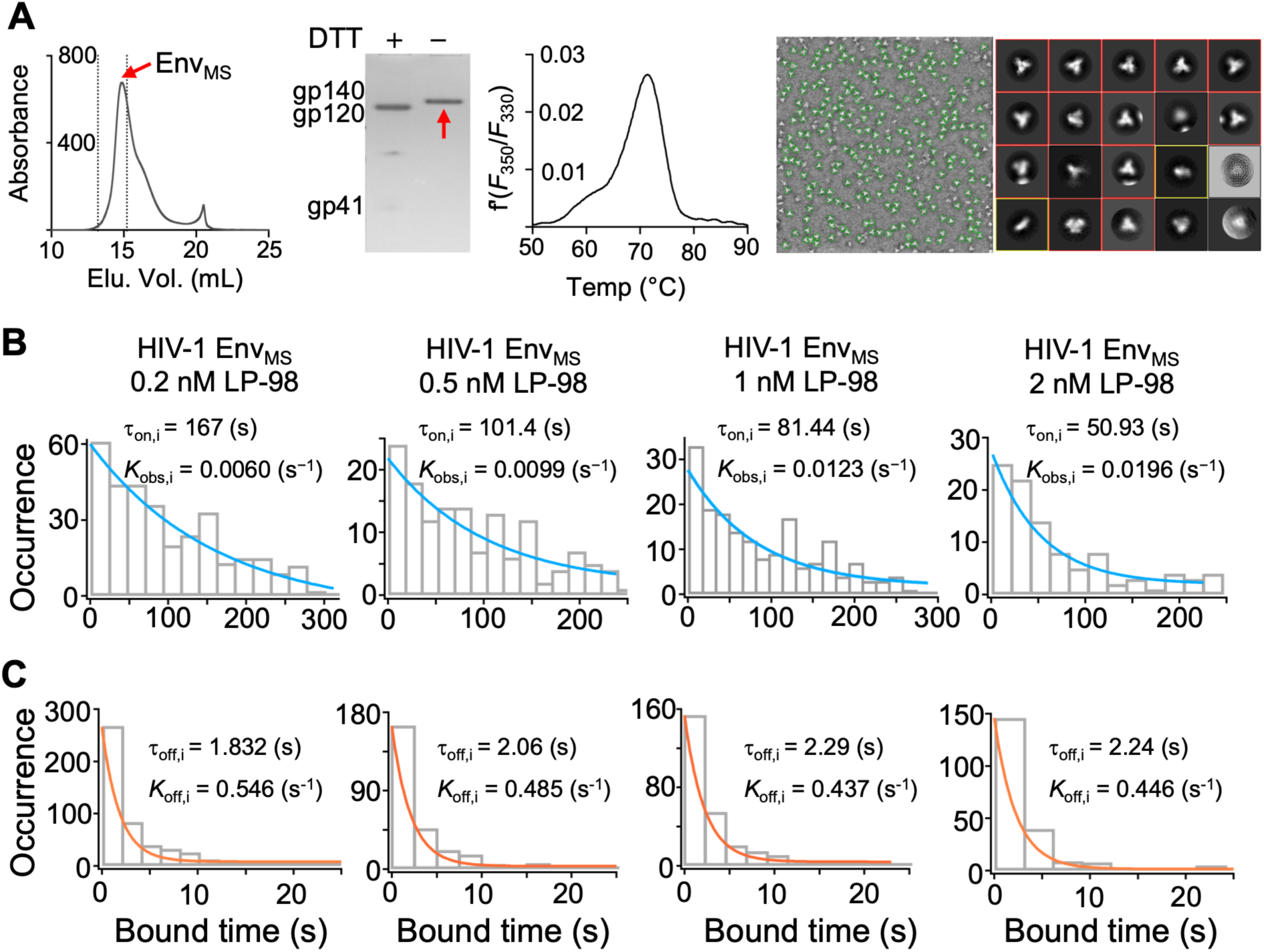
Single-molecule quantification of LP-98 binding kinetics to Env_MS_. **(A)** Purification and characterization of the Env_MS_ by SEC, SDS–PAGE, DSF, and NSEM. **(B)** Histograms of unbound dwell times yield *K*_obs,i_ at 0.2, 0.5, 1, and 2 nM of LP-98 interacting with Env_MS_. **(C)** Histograms of unbound dwell times yield *K*_off_ of LP-98 interacting with Env_MS_.

**Figure S7.**
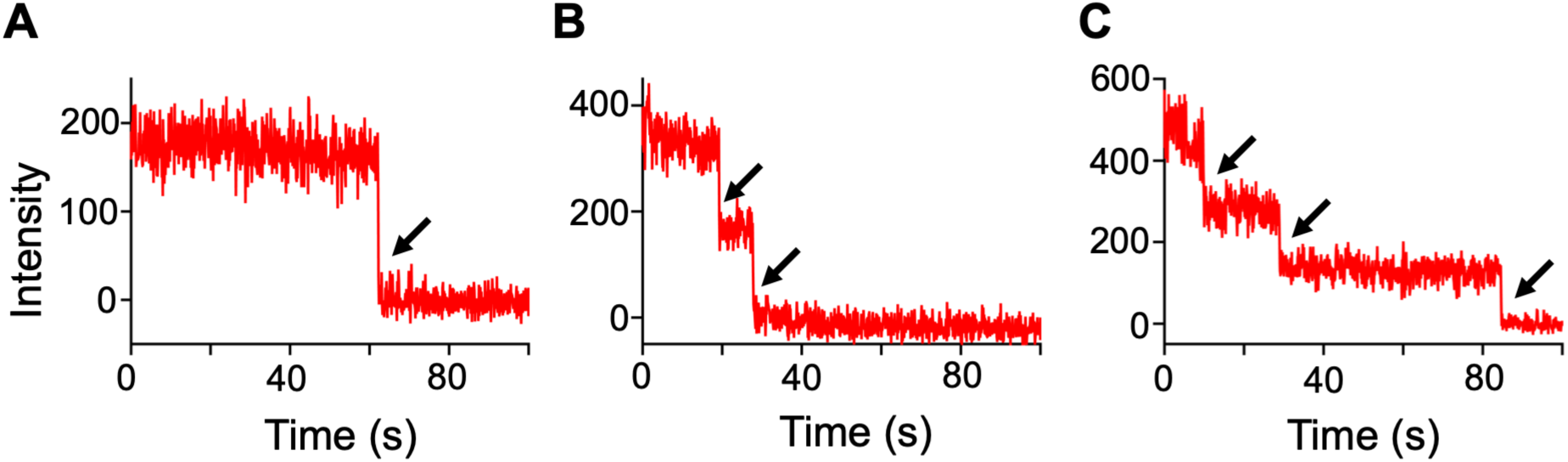
Stepwise photobleaching reveals the number of fluorescent labels per trimer. (**A-C**) Representative fluorescence trajectories of Env_SS_ show single-step (**A**) and stepwise photobleaching events (**B** and **C**), highlighted by arrows. These trajectories represent trimers containing one (A, the predominant labeling case), two (B), or three (C) fluorescent labels per trimer.

**Figure S8.**
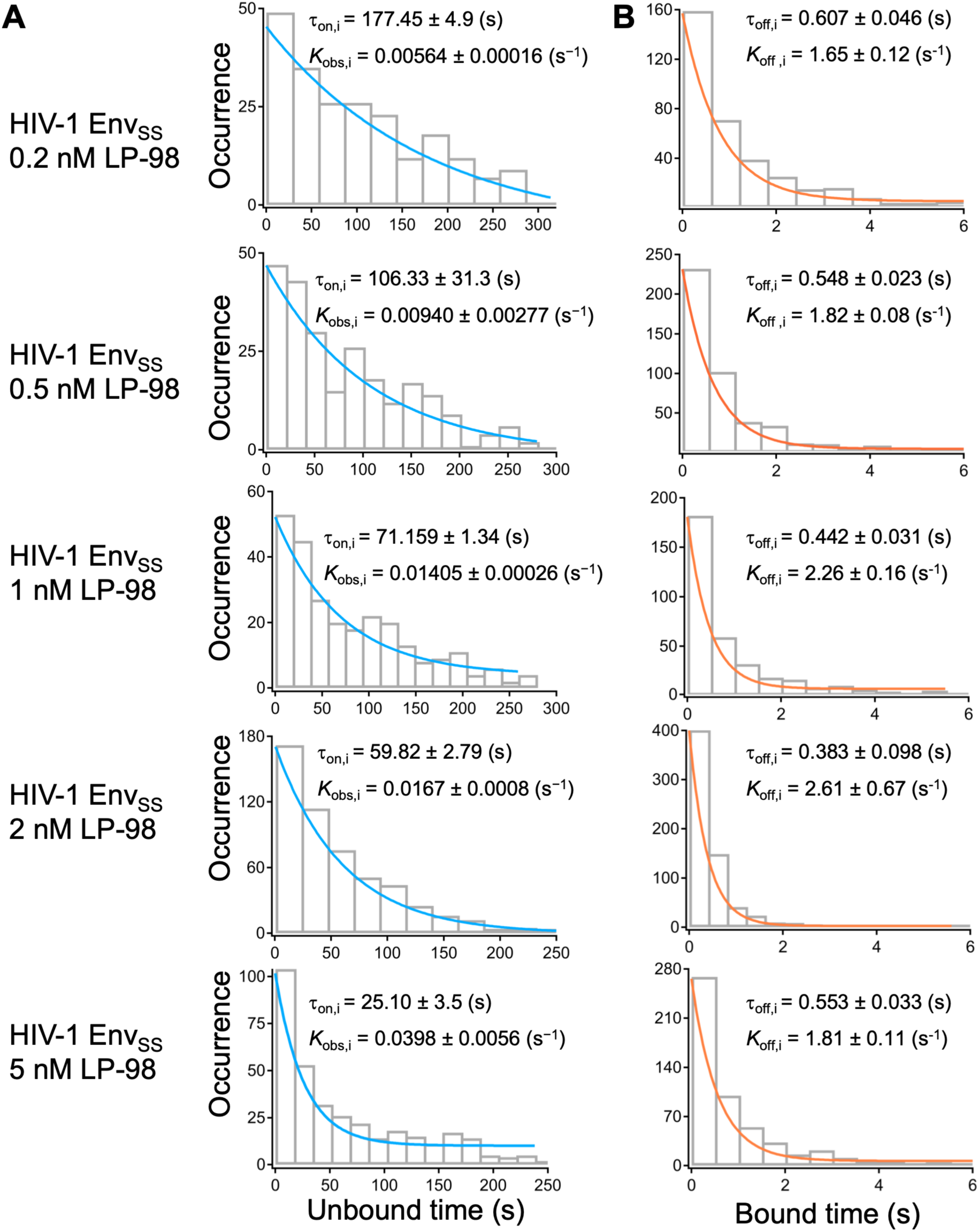
Unbound and bound dwell-time distributions of LP-98 on native-like prefusion Env_SS_ trimers. **(A)** Histograms of unbound dwell times yield *K*_obs,i_ at 0.2 nM, 0.5 nM, 1 nM, 2 nM, and 5 nM of LP-98 interacting with Env_SS_. **(B)** Histograms of bound dwell times yield *K*_off,i_ of LP-98 interacting with Env_SS_.

**Figure S9.**
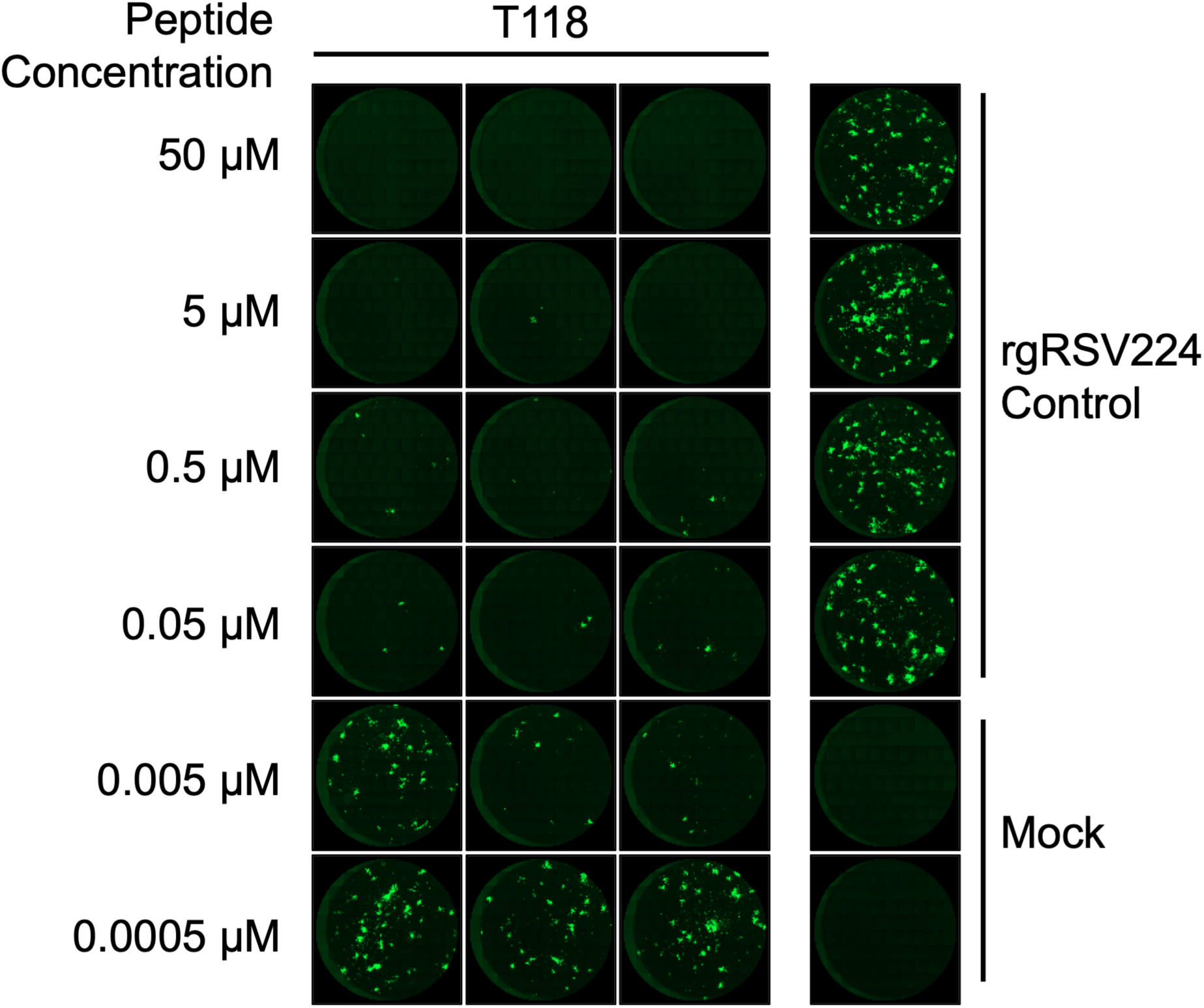
Inhibition of RSV infection by T-118 using a fluorescence plaque assay. Fluorescence images of HEp-2 cells infected with RSV carrying a GFP reporter in the presence of increasing concentrations of T-118 were used to quantify the inhibitory ability of T-118 against RSV (Fig. 4C) based on fluorescence intensity across the area of each image.

**Figure S10.**
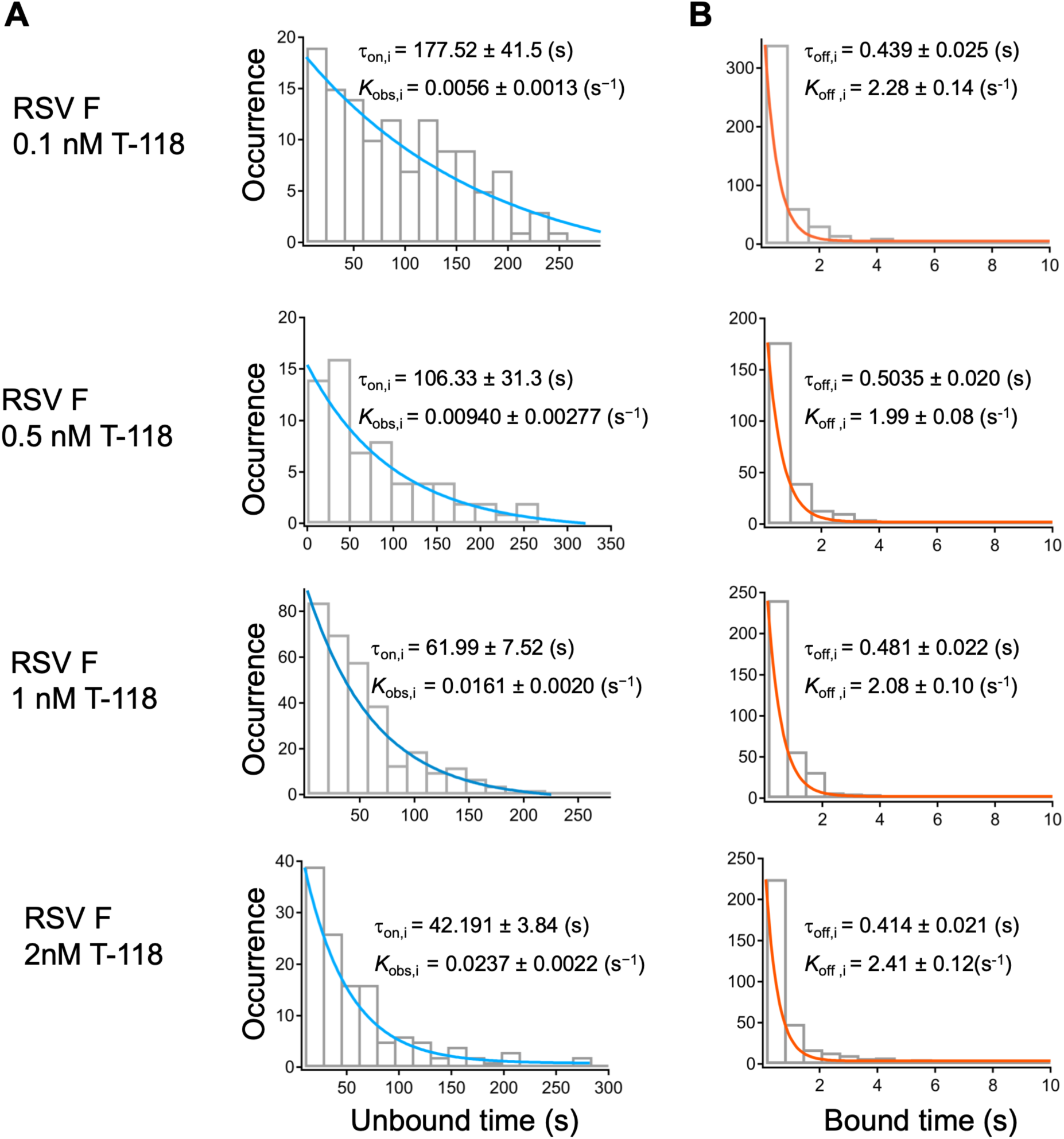
Unbound and bound dwell-time distributions of T-118 on RSV native-like prefusion F trimers. **(A)** Histograms of unbound dwell times yield *K*_obs_ at 0.1 nM, 0.5 nM, 1 nM, and 2 nM of T-118 interacting with prefusion F trimers. **(B)** Histograms of bound dwell times yield *K*_off_ of T-118 interacting with prefusion F trimers.

**Figure S11.**
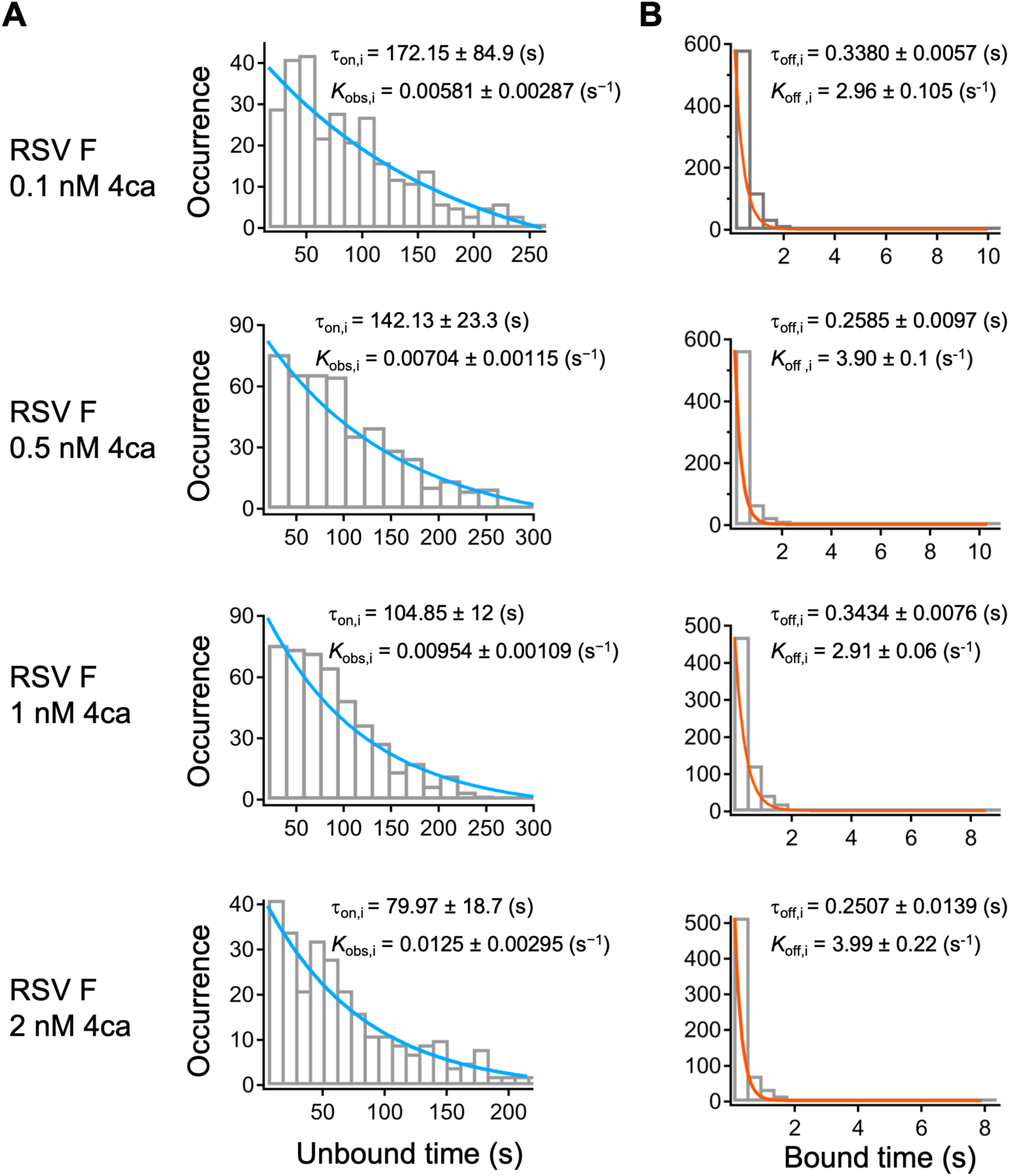
Unbound and bound dwell-time distributions of 4ca on RSV prefusion F trimers. **(A)** Histograms of unbound dwell times yield *K*_obs,i_ at 0.2 nM, 0.5 nM, 1 nM, and 2 nM of 4ca interacting with prefusion F trimers. **(B)** Histograms of bound dwell times yield *K*_off,i_ of 4ca interacting with prefusion F trimers.

